# Natural selection driven by escape from shifting antibody classes shapes SARS-CoV-2 evolution

**DOI:** 10.64898/2026.03.19.712895

**Authors:** Charlie Hamilton, Mahan Ghafari, Alice Ledda, Katrina Lythgoe, Christophe Fraser, Luca Ferretti

## Abstract

The phenotypic fitness landscape defines the action of natural selection on pathogens, linking changes in their phenotypes to transmission and evolution. The rapidly changing nature of epidemic spread and antigenic landscapes pushes viruses to evolve on fitness seascapes. As a result, evolution of viruses such as SARS-CoV-2 proceeds in a neverending series of waves, driven by epistatic interactions and by the arms race between viral adaptation and human immunity. Phenotypic characterisation of these rapidly changing fitness seascapes is an open challenge. Using a Phenotypic Selection Inference framework that links phylogenetic estimates of mutation fitness effects with deep mutational scanning data, we traced how selective pressures on viral phenotypes have shifted throughout the COVID-19 pandemic. Natural selection has favoured enhanced ACE2 binding since the emergence of SARS-COV-2, with relatively constant selective pressure even for the most recent variants. The strength of selection for antibody escape was comparable to ACE2 binding during early evolution, but as population immunity rose, escape from class 3 and then class 2 antibodies became dominant. For variants circulating in 2024, natural selection shifted toward class 3 antibody escape, while those circulating in 2025 have experienced dynamic, rapidly changing pressures for escape from all antibody classes. These transitions reflect an ongoing arms race between viral adaptation and human immunity. Our findings reveal that SARS-CoV-2 antigenic evolution is governed by dynamic, class-specific immune pressures, and that selection for replication capacity has been continuously present during the pandemic, presumably to compensate for the effects of antigenic escape on viral replication. Our approach for the inference of phenotypic selection provides a framework to understand and anticipate the evolution of future variants.

## Introduction

SARS-CoV-2 evolved at a remarkable speed both molecularly and antigenically ever since its emergence in 2020. These evolutionary changes have been widely studied both per se, with many works examining mutation rates, recurrent mutations, and their possible fitness ^1–5^ and in terms of the changing phenotypic traits of the virus, such as the appearance of major variants with enhanced intrinsic transmissibility (e.g. Alpha) or immune escape (e.g. Omicron)^6–8^. The antigenic evolution of SARS-CoV-2 has been studied in detail by measuring neutralisation of many strains by many antibodies/sera^9,10^ and through structural biology ^11–13^.

A glimpse into the most relevant part of the genotype-phenotype map for SARS-CoV-2 has been obtained through deep mutational scanning (DMS) experiments^14–16^, high-throughput experiments that determine the phenotypic effect of a large range of single amino acid changes (mutations) to a particular sequence of a major virus lineage, by creating a library of mutated sequence copies. These experiments have been mostly confined to the Spike protein or its receptor binding domain (RBD).

Unfortunately, merely observing evolutionary changes, both genetic and phenotypic, is not sufficient to understand their drivers. Many evolutionary changes are driven by natural selection, which favours individual viral sequences whose phenotype has a higher fitness than the rest of the population. Exactly which evolutionary change will occur at a specific time point depends both on the fitness impact of the change - i.e. the local fitness landscape - and the composition of the population. The action of natural selection alters the genetic composition of the population over the (viral) generations, shifting it towards regions of higher fitness. Reconstructing selective pressures from these observed evolutionary changes is an inverse problem that requires specific approaches. Several such approaches have been developed to examine the fitness of individual SARS-CoV-2 mutations ^4,1,17,18^. Among these methods, state-of-the-art phylogenetic approaches allow us to infer the fitness effects of almost all amino acid mutations across the whole SARS-CoV-2 genome from their frequency and pattern of appearance along the phylogenetic tree^4^.

Selection acts directly on phenotypes, rather than genotypes. Phenotypic traits of SARS-CoV-2 that may have been under natural selection include the components of the virus’s intrinsic transmissibility, namely its ability to bind to ACE2 cell surface receptors and successfully enter, replicate within, and then exit the cell. Another central set of phenotypic traits concern the ability of the virus to escape immune defences, especially by preventing neutralizing antibodies from binding to it. Antibody escape can be further broken down into the kind of antibodies that are escaped, classified by their target epitope. One example classification is the four anti-RBD Barnes classes^19^, determined by which conformations of the RBD it binds to and whether the antibody blocks ACE2 binding. Detecting selection on all these phenotypes is key to reconstruct the evolutionary history of the pathogen and to predict the properties of future variants.

It is expected that the strength of selective pressure for different phenotypes has changed since the emergence of SARS-CoV-2. Shifts in selective pressures can be caused by changes in the underlying fitness landscape of the virus over time, which is known as a fitness seascape ^20^. For example, the viral fitness landscape is strongly linked to the antigenic landscape of the host population. Mutations that escape a particular antibody will only be beneficial if (i) the antibody is widely present in the population due to immunity from previous infections/vaccination, such that escaping the antibody increases the susceptible population; (ii) that antibody has a strong affinity for the currently circulating variant of the virus: if novel strains emerge with different antigenic profiles and low affinity for the antibody, escape mutations in these strains confer no advantage. We expect that shortly after emergence of a new highly infectious pathogen in a fully naive population, the main selective pressure would be towards increasing transmissibility among humans. However, after the pathogen has infected most of the population, natural selection would instead favour variants able to escape the existing immunity. An apparent fitness seascape can also be caused by changes in the viral genetic background that affect the local fitness landscape (epistasis). For all these reasons, fitness landscapes are often both time- and environment-dependent.

Our aim is to determine which phenotypes have been under natural selection and the strength of these selective pressures over time. We exploit the fact that the genetic fitness landscape is a combination of genotype-phenotype map and the phenotypic fitness landscape. Or, equivalently, the fitness effect of a mutation results from a combination of the phenotypic effects of the mutation and the selective pressure on these phenotypes. Decomposing time-dependent mutational fitness effects inferred through phylogenetics and phenotypic effects measured by DMS studies, we reconstruct for the first time the fitness seascape of SARS-CoV-2, exploring the selective pressures acting on the viral phenotypes since its emergence and how they predict the properties of emerging variants.

## Results

### Post-Omicron selection for escape from shifting antibody classes

The novelty of this work lies in reconstructing the phenotype–fitness landscape, i.e., the selective pressures acting on viral phenotypes and how they change over time. The long-term genetic and phenotypic evolution of SARS-CoV-2 has proceeded in saltatory shifts arising from the stochastic emergence of major variants ^8^. In this section, we analyse how this landscape has changed across successive variants.

We develop a new approach for “Phenotypic Selection Inference” to infer the contributions of different phenotypes to the fitness landscape, and therefore estimating the selective pressure acting on them (Figure 1A). For each mutation in our analysis, its time- or clade-dependent fitness effect Δ*f*(*t*) is obtained from a phylogenetic approach, and the phenotype effects for *i*th trait relative to a specific genetic background Δϕ_*i*_ are obtained from DMS studies. We model the fitness effects Δ*f*(*t*) as a function of all phenotypic effects Δϕ_*i*_ across all mutations.

**Figure 1.**
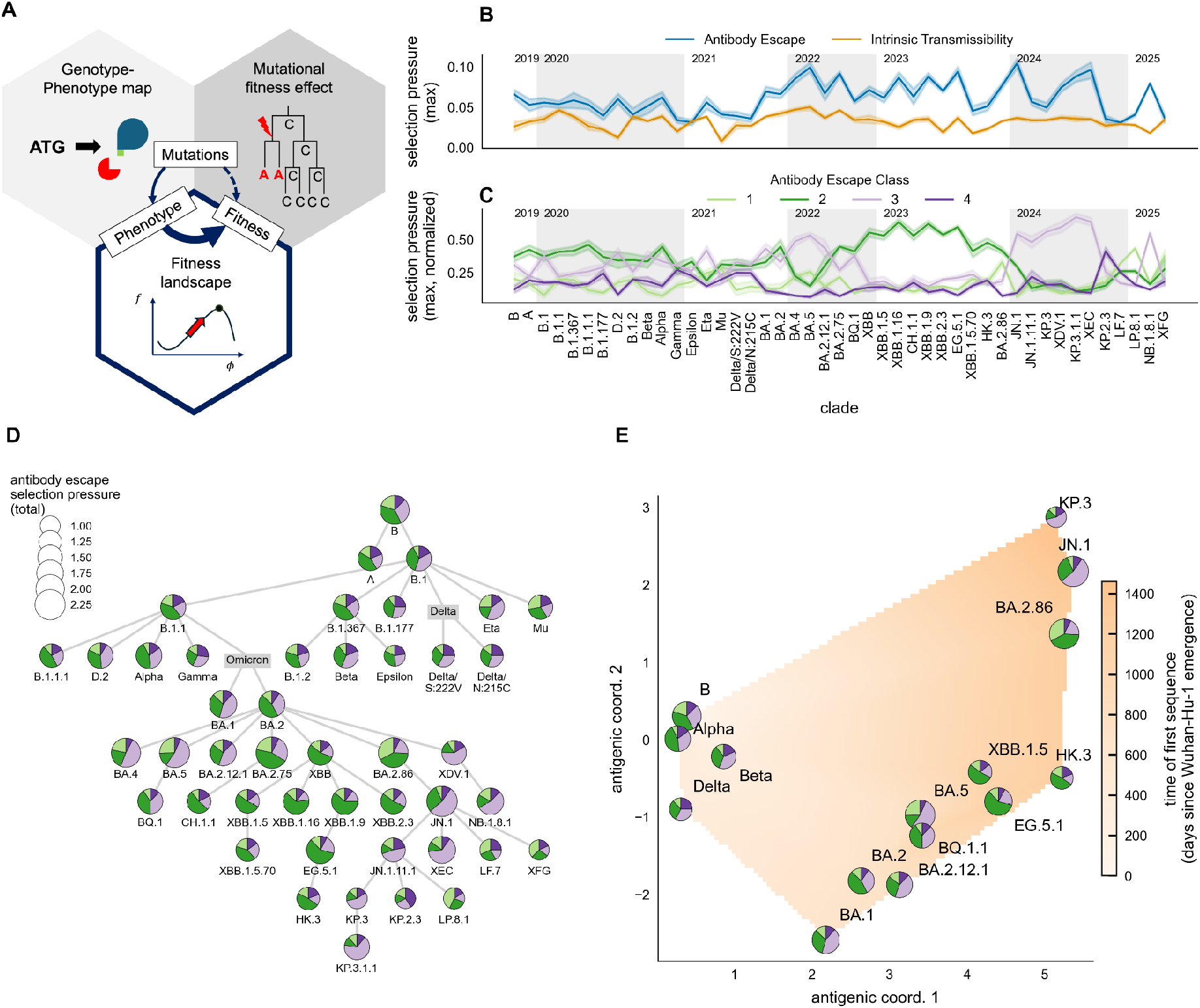
The fitness seascape of SARS-CoV-2. (A) The relationship of the phenotypic fitness landscape to the genotype-phenotype map and the genotype-fitness map. For mutations, the genotype-phenotype (GP) map links mutations with each of their associated phenotypic changes. The fitness landscape is the mapping between these phenotypes and viral fitness. The genotype-fitness map, also known as genetic fitness landscape, is the composite map (dashed arrow) of the GP map and the phenotypic fitness landscape. Knowing two of these arrows/relations allows us to determine the missing one. For this work, we leverage the GP map (from deep mutational scanning data) and the genotype-fitness relation (from the phylogenetic estimator of mutational fitness effect) to determine the missing side of the triangle, i.e. the phenotypic fitness landscape. (B) The transition between intrinsic transmissibility and immune (antibody) escape. Plotted for each Nextstrain clade is the maximum selective pressure for phenotypes within each of the two classes. Around the emergence of Omicron BA.1 we observe a large increase in the maximum selective pressure for antibody escape. (C) Transitions in the selective pressures for escape from different antibody epitope classes. Plotted are the maximum selective pressure across antibodies in each of the four Barnes anti-RBD epitope classes, normalised to sum to unity for each major variant. We observe clear transitions in the dominant escape epitope pressure. The same data is embedded in phylogenetic space (D) and antigenic space (E). Variants are placed in antigenic space using existing antigenic cartography data from Rössler et al.^43^. The antigenic space is coloured by time of emergence of the variants.

For small changes of phenotype ϕ − ϕ^*wt*^ compared to a “wild-type” virus, the fitness landscape *f*(ϕ) can be approximated as a linear function of the changes in phenotype:

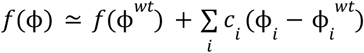

The coefficients *c*_*i*_ quantify the contribution of a particular phenotype *i* to the fitness, i.e. the strength of selection (‘selective pressure’) on that phenotype. Our approach infers these coefficients by decomposing this phenotype-fitness relation in terms of mutational effects of a mutation µ:

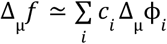

where the phenotypic (or fitness) effect Δ_µ_ϕ of a particular mutation µ is given by the difference between the value of the phenotypic trait (or fitness) in viruses with and without the particular mutation.

Mutational fitness effects are inferred from public mutation-annotated phylogenetic trees ^21^, using the method from Bloom and Neher^4^, based on the the relation between fitness effect Δ*f* of a particular mutation and its rate of appearance in the tree. Counts of mutations, and hence fitness effects, will vary across the phylogenetic tree; in our analyses we estimate the fitness within specific Nextstrain clades^22^, and/or within specific time windows. Phenotypic effects of mutations are taken from existing deep mutational scanning (DMS) data (compiled in Greaney et al.^23^ and Taylor and Starr^24^).

We focus on two different sets of phenotypes: antibody escape, and transmissibility (i.e. ACE2 binding). We use two different reference (‘wild-type’) sequences for each of the two sets: for transmissibility, the founder sequence of a major clade of interest, and for antibody escape the Wuhan-Hu-1 sequence. As in general there will be substantial covariance between different phenotypes, we use ridge regression to penalize the size of regression coefficients, and account for the collinearity. We standardise all phenotype effects to have a mean of zero and unit variance across mutations, interpreting the standard coefficient from the regression as the phenotypic contribution to fitness, and hence the selective pressure (Figure 1A).

We compare the strength of the selective pressures from antibody escape and intrinsic transmissibility over different clades ordered by their time of emergence. The resulting seascape is shown in Figure 1B. We compare clades by the maximum absolute standardized coefficient across phenotypes, a statistic that allows group-level comparison of selection strength independent of group size. We find a clear selective pressure for increased ACE2 binding across major variants and over time (Figure 1B, blue line). Selective pressure for antibody escape is present from the beginning and closely tracks the pressure on transmissibility until a transition occurs with the appearance of Omicron. After the appearance of Omicron, which also corresponds to the time of a large drop in naive individuals, selective pressure for antibody escape increases significantly. On the other hand, the strength of selection for increased transmissibility remains approximately constant until 2025 (Figure 1B, orange line).

Zooming in on the escape pressure, we explore the variation in the maximum selective pressure among antibodies that belong to a particular epitope class (Figure 1C), namely the four Barnes epitope classes^19^ defined by which conformations of the RBD the antibody binds to and whether the antibody blocks ACE2 binding. After the emergence of Omicron and the increase in population immunity due to infection and vaccination, immune escape has had an elevated contribution to natural selection acting on SARS-CoV-2 (Figure 1C). This drives the dynamics of selective pressure and immune escape for individual antibodies.

The strength of the fitness contribution for different antibody epitope classes shows a marked temporal dynamics. This is clear from the variation between clades arranged in time order (Figure 1C) as well as from their distribution in phylogenetic (Figure 1D) and antigenic space (Figure 1E). For antibodies in the same epitope class there are considerable similarities between the time variation of the phenotype’s contribution to overall fitness (Supplementary Figure S4), suggesting there are generalised selective pressures broadly shared by multiple antibodies, and may be as broad as a particular Barnes epitope class.

Class 2 antibody escape contributed the most to viral fitness throughout until BA.1. After the appearance of Omicron BA.1, the importance of class 1 and class 3 antibodies has increased. For some BA.2 descendants that appeared independently shortly after in 2022 (BA.4, BA.5, etc), the dominating selective pressure was for class 3 escape. The recent trends are especially interesting. All recent variants are descendants of BA.2, which was dominated by selective pressure for escape from class 2 antibodies. Among the recent variants, the immediate descendant of BA.2 was BA.2.86, which experienced a shift towards class 1 escape. Successive variants such as KP.3 saw a collapse in the contribution of class 1 escape and, more remarkably, of class 2 escape as well. For the first time, for these variants circulating in 2024, natural selection is driven by pressure for escape from class 3 antibodies, with a negligible role for classes 1 and 2. Finally, in the latest twist, selection on recent variants circulating in 2025 has been alternating between all four antibody classes, with the latest currently circulating variant XFG showing another shift towards escape from class 2 antibodies.

### Selective pressures change over short timescales

Changes in selection occur across multiple time scales. While we have described how selection changed across different clades, these pressures are likely to change even within a single clade from one week to the next. For selected SARS-CoV-2 variants, we examine the changes in selective pressure within a specific variant through the variation of the contribution of the phenotype to fitness for different phenotypes. We restrict this analysis to sequences from England, as epidemiological time trends often differ geographically (see Methods for more details).

We observe non-trivial changes in selective pressures over time within different clades. For example, for Omicron BA.5 (Nextstrain clade 22B) we explore the linear trends of increase/decrease in selection strength over time across many different phenotypes (Figure 2, bottom left). We observe a range of trends for selection strength, from rapid increase for class 3 antibody escape to rapid decrease for class 2. The explanation lies in the peculiar nature of selection for immune escape in BA.5. Its patterns of natural selection are unusual, because of a strong pressure for escape from class 3 antibodies, contrasting with a much weaker pressure to escape class 2 antibodies – a behaviour not observed in other Omicron variants, with the partial exception of the contemporaneous variants BA.4 and BA.2.12.1. Our results show that the shift in pressure from class 2 to class 3 antibody escape was not concentrated at the time of emergence of BA.5, but it occurred gradually, becoming progressively more pronounced during the whole BA.5 wave.

**Figure 2.**
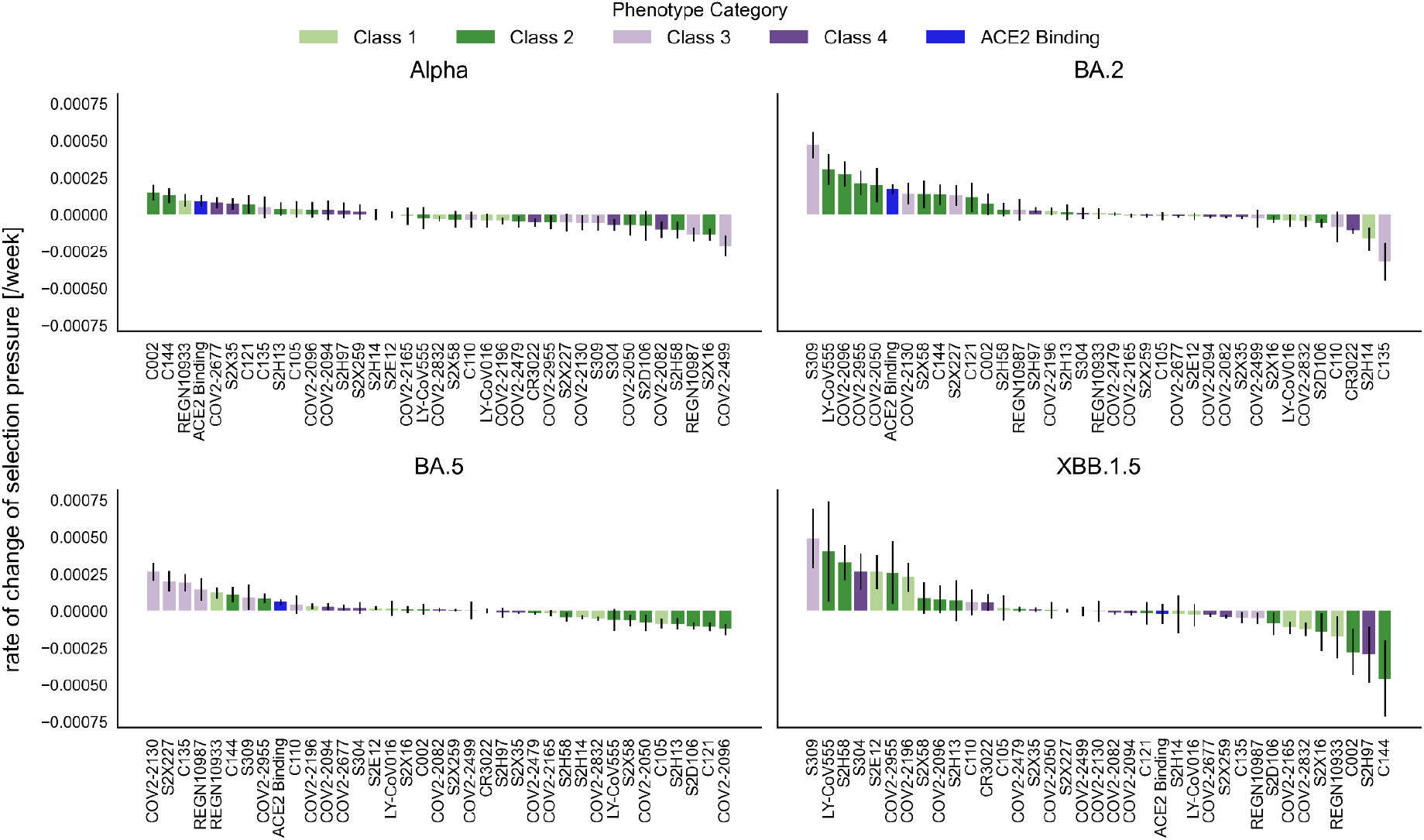
Short-time trends in selective pressures within clades. Rate of change of the selective pressure with time (per week) for different phenotypes, as measured within different major variants (Alpha, BA.2, BA.5, XBB.1.5). The bars for each phenotype are ordered by the value of the selective pressure, and coloured according to the class of phenotype they belong to.

The picture differs significantly for other variants (Figure 2). Within Alpha, selection decreases in time for a majority of antibodies. BA.2 shows the opposite pattern, with a strong increase in selection for escape from class 2 and 3 antibodies, likely reflecting the transition to increased pressure for immune escape for early Omicron variants. XBB.1.5 shows a mixed pattern, with examples of increasing positive selection for escape from antibodies from all four classes. Interestingly, selection for improved ACE2 binding increases significantly in time within all variants except XBB.1.5, for which it remains approximately constant. This trend is at odds with expectations of pressure for compensatory evolution soon after the emergence of a new variant.

### Selective pressures predict within-clade evolution

Understanding selective pressures opens new opportunities to predict short-term evolution. Evolutionary changes (in both genotype and phenotype) are the result of natural selection acting on the variation within the current population. Since the relation between selection and evolution is mediated by the genetic composition of the population, knowledge of this composition is needed for short-term predictions.

The dynamics of individual mutations provide a clear illustration of the relationship between natural selection - summed up in the virus fitness, *f* - and evolutionary change - i.e. the time variation in the frequency *x* of a mutation circulating in the viral population. Changes in the population frequencies of mutations are caused by natural selection, and their rate of change is given by the equation 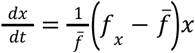 where *f*_*x*_ is the average fitness of viruses containing the mutation, and 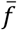 is the average fitness of the population. The term 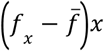 is the population-level covariance between viral fitness *f* and presence/absence of the mutation. It depends not only on the fitness effect of the mutation, but on how the mutation is distributed within the population, i.e. its frequency and its association with other beneficial or deleterious mutations. Hence, it will change in time as the genetic composition of the population changes.

This covariance can be immediately used to predict short-term evolution. In Supplementary Figure S24, we illustrate how for some examples of mutations in the SARS-CoV-2 Spike protein, evolutionary changes - i.e. the overall change in mutation frequency - are tracked remarkably well by the cumulative covariance, when the latter is computed using fitness effects inferred by phylogenetic methods^4^.

Natural selection acts directly on phenotypes, rather than individual mutations, and is described by the phenotypic fitness landscape of the virus, that is, a map from phenotypes to viral fitness. The general formalism for analysing the link between fitness, phenotype, genetic variation and selection was first derived by Price ^25,26^. Assuming perfect heritability and no transmission bias, the classical Price’s equation for phenotype evolution reduces to 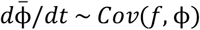 where the covariance between fitness and phenotype *Cov*(*f*, ϕ) is calculated across all viruses in the population. Based on this formalism, the population-level covariance between the fitness *f* and a phenotype is the best predictor of its short-term evolution for the phenotype.

Figure 3 shows how natural selection for a particular SARS-CoV-2 phenotype, as encoded in the phenotype’s fitness landscape, induces population-level shifts in the average value of the phenotype. Here we compute phenotype and fitness from viral sequences, assuming that both are well approximated by a linear combination of the effects of the associated mutations. For example, the fitness of Omicron BA.5 viruses correlates with the escape from a monoclonal antibody (Cilgavimab/COV2-2130), that is, viral variants that are able to escape Cilgavimab are able to infect more hosts and hence favoured by natural selection. The Cilgavimab-escape variants that are present in the population may harbour different combinations of mutations, but they all become more abundant under selective pressure on this escape phenotype. The average value of Cilgavimab escape in the viral population 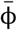 therefore increases in time, again following Price’s equation.

**Figure 3.**
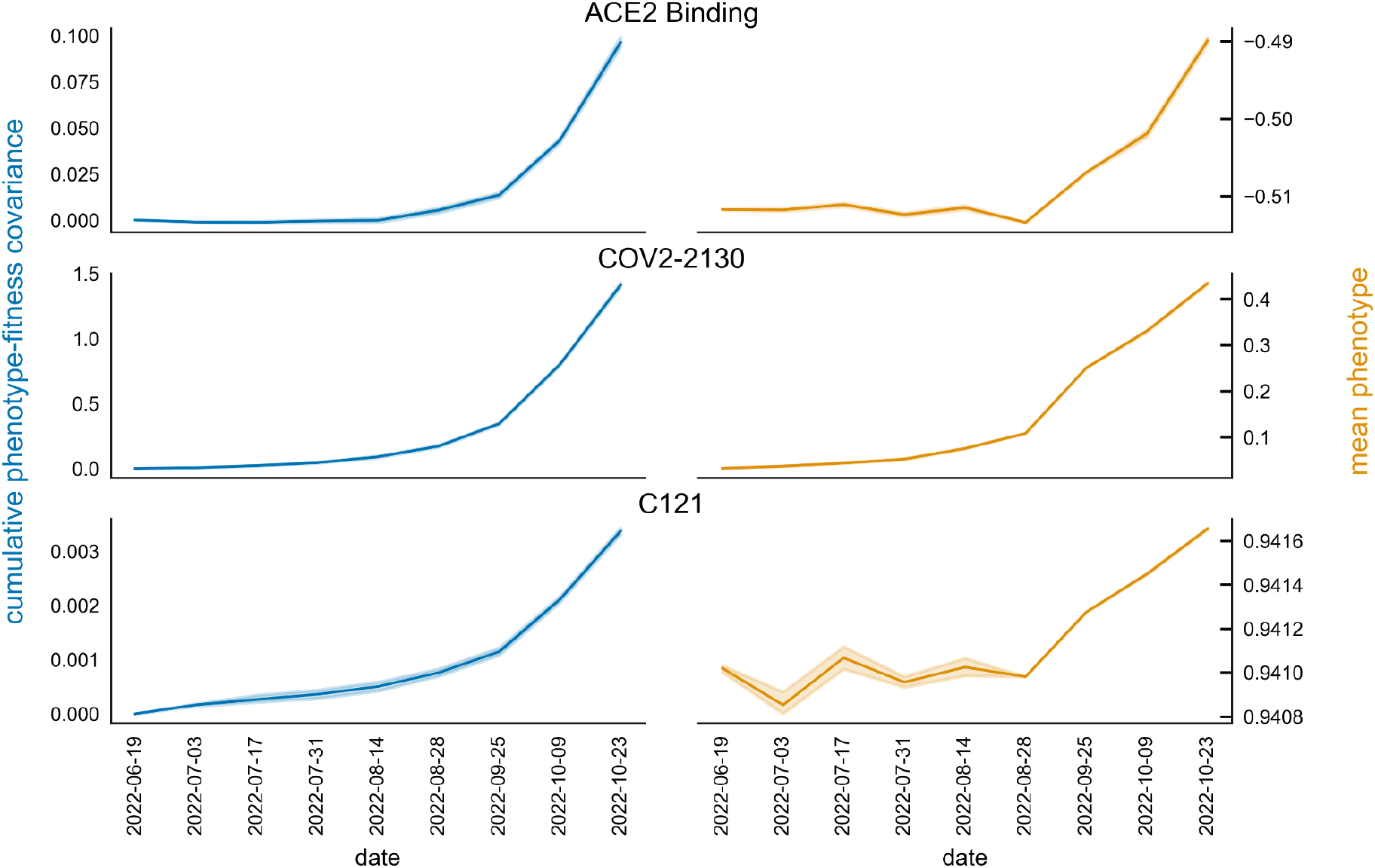
Predicting short-term evolution of SARS-CoV-2 within the BA.5 clade. The estimated covariance between fitness (inferred by phylogenetic approaches) and phenotype (inferred by DMS studies) across viral sequences can be used to predict phenotype evolution within a clade. In fact, the cumulative value in time of the estimated covariance (left) and the average value of the phenotype over viral sequences (right) increase approximately proportionally with time as predicted by Price’s equation, since the change in the average phenotype 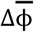 is proportional to the true population-level covariance *Cov*(*f*, ϕ)).

This result holds across all phenotypes ϕ, i.e. their time evolution within a clade closely follows the cumulative covariance *Cov*(*f*, ϕ) as predicted by Price’s equation (Supplementary Figure S25). This confirms that the phylogenetic fitness measurements capture the actual fitness landscape and that Price’s equation can be used for short-term evolutionary predictions.

### Selection drives evolution of antibody escape

Selective pressures not only shape short-term evolutionary dynamics, but can also anticipate long-term evolutionary changes. To better understand the temporal relationship between selection and SARS-CoV-2 evolution, we introduce a conceptual framework for the dynamics of immune escape from an individual antibody.

As an antibody grows in prevalence in the human antigenic landscape, the selective pressure to escape it grows proportionally to its population-level prevalence. Consequently, escape variants are rapidly favoured and the circulating viruses tend to be less and less neutralised by the antibody^27^. However, this effect slows down because of diminishing returns, which make the escape from a specific antibody less and less relevant for the fitness of the virus once most circulating viral sequences are already able to escape the antibody. Such diminishing returns are a common feature of fitness landscapes^28^, where the rate of fitness advantage often decreases with increasing fitness: for example, the first escape mutations cause a much bigger loss of neutralisation than later additional escape mutations. They can also be caused by negative frequency-dependent selection, due to a combination of cross-neutralisation and immune waning^29^, resulting in a decreased fitness for a given strain once a significant part of the population has been infected with the strain. Diminishing returns imply that selection for escape from a specific antibody is expected to show a characteristic bell-like dynamics, where the ability of the antibody to neutralise the virus is low before the selection peak, and high after the peak. Such dynamics can be quantitatively recovered from simple SIR-like models^30^ (see Figure 4B).

**Figure 4.**
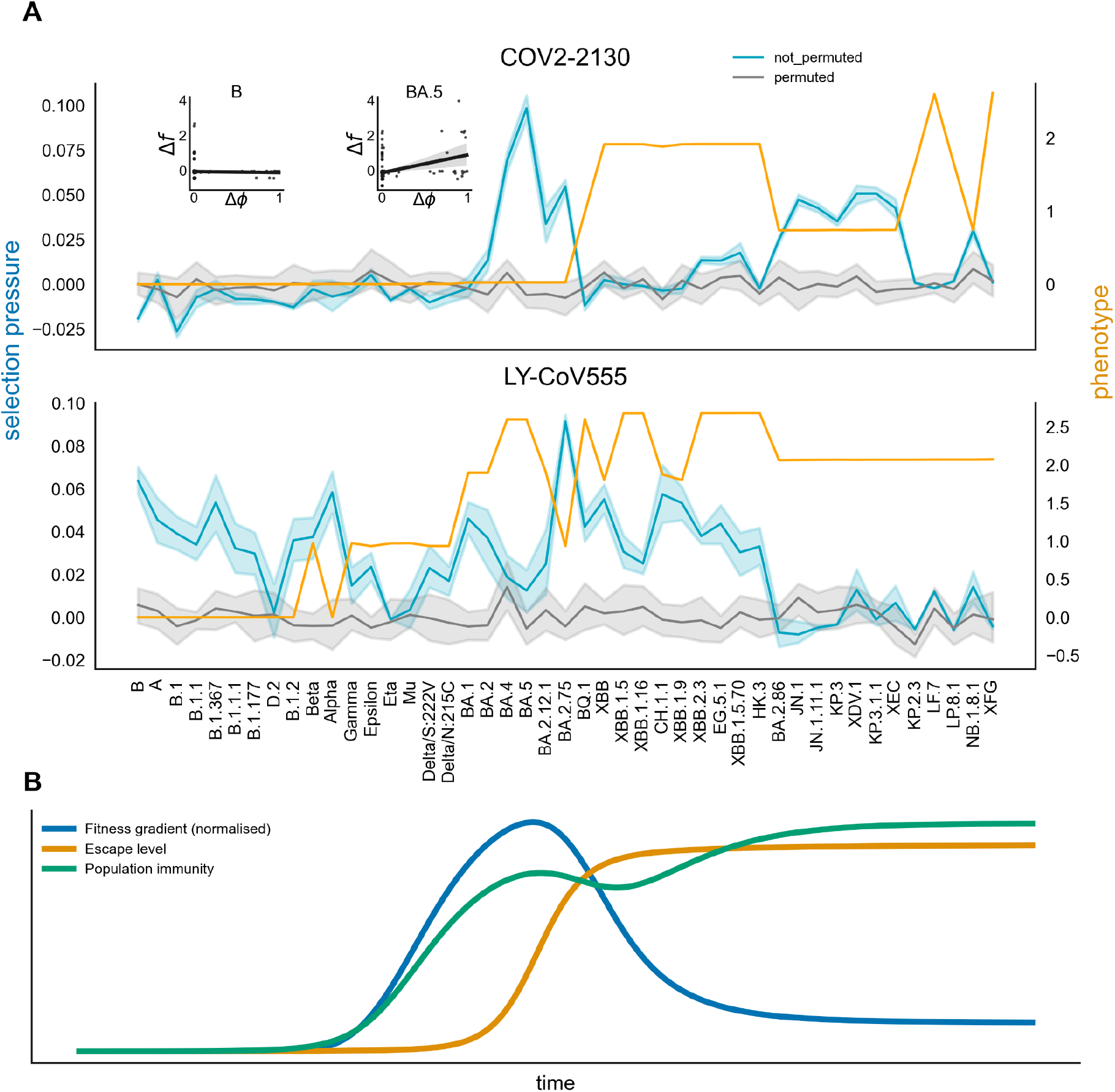
Dynamics of selective pressure and escape from an individual antibody. (A) The selective pressure (phenotypic contribution to fitness, blue) for escape from the antibody COV2-2130 (Top), and the realized phenotype (escape fraction, orange) for major clades/variants ordered in time. The phenotype of a particular variant is given by the sum of the DMS phenotypic effects of the defining mutations. A bump in selective pressure for escape for BA.4/5 precedes an increase in the realized escape from the antibody. In insets are shown the fitness landscape (each marker is a mutation) for the major variants B (Nextstrain 19A) and Omicron BA.5 (Nextstrain 22B). Bottom: as above, but for a different antibody Ly-CoV555. (B) Dynamics of selection for escape from a generic antibody, from a multi-strain SIR model. The population immunity (green) is the proportion of individuals recovered from the original strain. The fitness gradient (blue) is the average fitness difference between successive strains, weighted by their prevalence in the population of infected individuals. The escape level (orange) is the average escape fraction against the original strain among the infected individuals. Rising population immunity (antibody prevalence) causes a concomitant rise in the fitness gradient for escape from the antibody, and a subsequent rise in the average value of escape in the infected population.

The observed dynamics of escape from several antibodies follow precisely the diminishing returns behaviour discussed above (Figure 4A). The selective pressure for the monoclonal antibody Cilgavimab/COV2-2130 across different clades follows a bell-like trend when ordered by time of emergence (blue line, Figure 4A, top). We can compare this trend with the level of antibody escape for each clade (orange line, Figure 4A, top), as given by the sum of the escape effects of its defining mutations. Following the peak in the selective pressure, the realised escape increases up to a stable value where it remains for some time. Such dynamics can be interpreted as the result of strong natural selective pressure after the antibody becomes part of the immune response of already-infected individuals, due to a change in the population antigenic landscape or an epistatic shift. The presence of a particular antibody in the immune response ignites a selective pressure to escape it, followed by increasing population-level escape from the antibody, and finally weaker selective pressure due to diminishing returns, until some variants escape the antibody completely. Similar trends are observed for other antibodies (Supplementary Figure S7). In the longer term, viral escape may decrease again, e.g. due to changes in the host antigenic landscape or in the genetic background or the virus, or changing antigenicity as it evolves against more recent selective pressures. This may cause new pressure to ‘re-evolve’ the escape, as was the case for Cilgavimab after BA.2.86 (Figure 4A, top).

### Forecasting future variants’ phenotypes from current selective pressures

The long-term evolution of SARS-CoV-2 is shaped by the recurring emergence of major variants with distinct phenotypes^8^, arising primarily from persistent infections^8,31^. Variants that happen to have a strong selective advantage compared to existing strains – i.e. that spread more effectively – emerge and replace them^6^. Natural selection therefore acts as a sieve, filtering which variants successfully emerge and spread. Although the appearance of new variants is highly stochastic, after their appearance they are likely subject to the same selective pressures as any other competing lineage. This implies that recent selective trends on a given phenotype should broadly predict future phenotypic change, albeit with considerable uncertainty.

To test this idea, we develop a model to predict the phenotype of newly emerging variants. Predicting when a new variant will appear is unrealistic. We focus on variants once they have already emerged: we compare their phenotype to a recent baseline and relate the phenotypic change to the selective pressures acting on circulating viruses just prior to their emergence. We use a two-step logistic-linear regression model (see Methods). The logistic component accounts for the stochastic nature of appearance, selection and emergence of major variants, while the linear component estimates the magnitude of phenotypic change.

The model effectively predicts the probability of emergence of new variants with substantial phenotypic change based only on past selection pressures. As shown in Figure 5A, both a model trained on the full dataset and its validation on a test set perform well, demonstrating that despite considerable stochasticity, selective pressures are strong predictors of major phenotypic shifts. The model also relates selection strength to the magnitude of phenotypic change, albeit with considerable variability, reflecting the role of selection as a “tinkerer” and a sieve on stochastically appearing variants, rather than a direct causal force.

**Figure 5.**
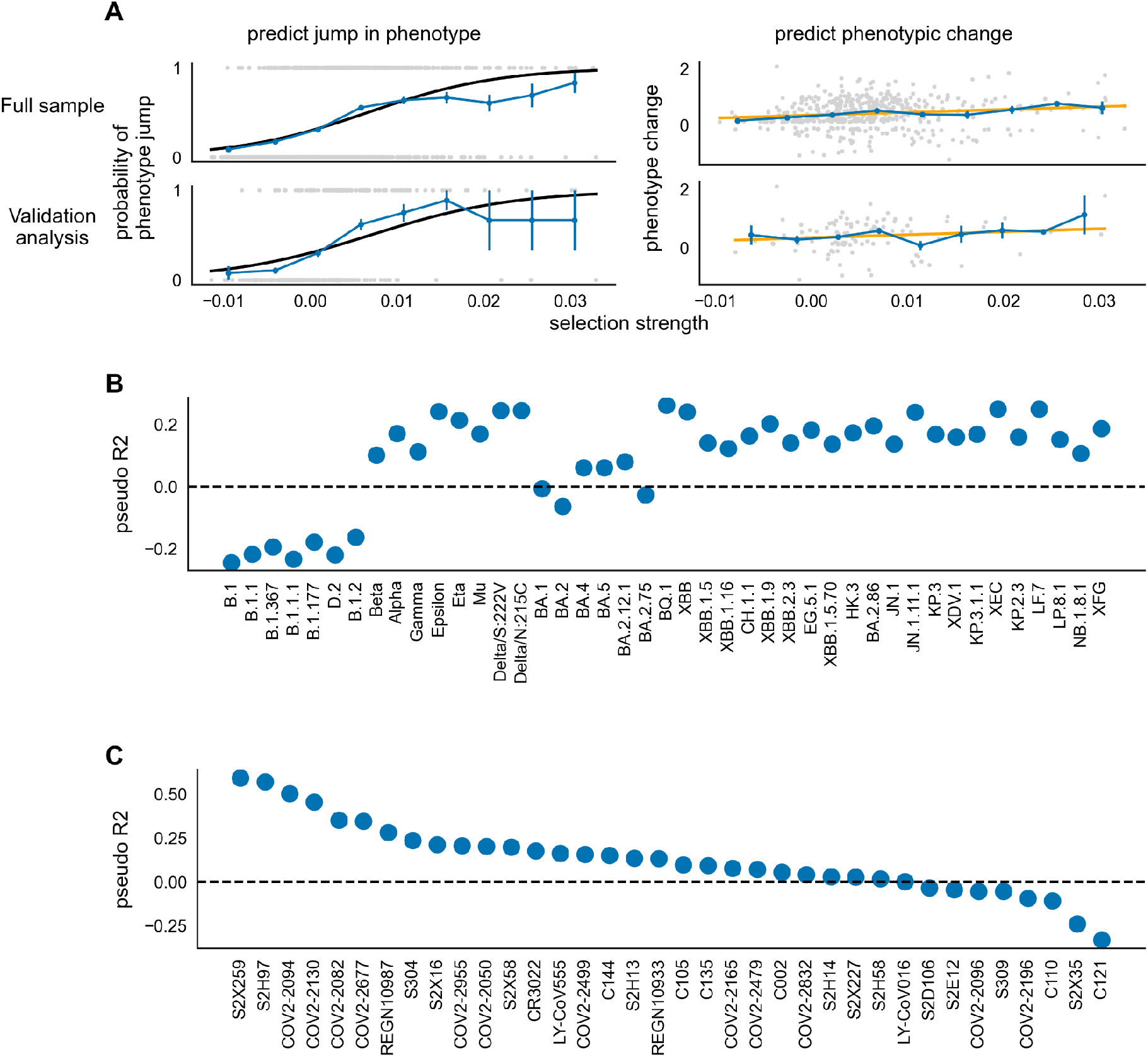
Prediction of phenotypic properties of new variants from recent selective pressures. (A) Predicting the emergence of major variants with substantial phenotype changes in terms of escape from an individual antibody. Left: empirical probability of substantive phenotypic changes (blue) as a function of selection strength on the phenotype, and corresponding fit from logistic regression (black). Right: empirical magnitude of substantive phenotypic changes (blue) as a function of selection strength on the phenotype, and corresponding fit from linear regression (orange). Predictive fits are shown for full-sample regressions (top row) and for out-of-sample validation using an 80/20 training/test split (bottom row). (B) Predictability of phenotype jumps for each variant, as estimated by McFadden pseudo-R2. This statistic is based on logistic versus null model fit to the full dataset. (C) Predictability of phenotype jumps for each phenotype, as estimated by McFadden pseudo-R2.

We examine how the predictability of substantial phenotypic shifts varied between different variants and phases of the pandemic (Figure 5B). Predictability would have been very low for the first variants at the beginning of the pandemic, since these variants show limited phenotypic changes. The evolution of the first major variants with a distinct phenotype such as Alpha, Beta, Gamma, Delta was predictable based on selective pressures, while the phenotypes of the first Omicron variants BA.1 and BA.2 would have been hard to predict, given the extreme evolutionary saltation event involved. Evolution of later Omicron variants is predicted by selective pressures at least until 2025.

Our model is reasonably predictive of the evolution of many phenotypes (Figure 5C). The easiest to predict are escape phenotypes under negative selection, for which the model correctly predicts that phenotypic jumps tend not to occur (since the wild type already shows no escape). Among phenotypes under positive selection, the evolution of escape from the monoclonal antibody Cilgavimab/COV2-2130 discussed in the previous section is the most predictable one based on previous selective pressures.

### Unpredictable patterns of antibody escape selection in 2025

Finally, we explore in detail the temporal dynamics of selection for the most recent SARS-CoV-2 variants, JN.1 (Nextstrain clade 24A) and its descent clades (Figure 6). While we observe a decreasing trend in time for selective pressure for class 2 and 4 escape (Figure 6C), the dynamics for individual antibodies are far from linear in time (Figure 6B). Class 2/3 escape pressures rise substantially but then fall again to zero. Class 1 escape pressure only rose around Autumn 2024. Class 4 is most important around Autumn/Winter 2024. These dynamics approximately reproduce the results of our clade-based analysis (Figure 6A). We note that the emergence of variant KP.2.3 (Nextrain clade 24G) appears to be coincident with the end of the long-time domination of class 3 antibody escape pressures (Figure 6A) that had lasted since the emergence of JN.1, to be replaced by a more chaotic regime in which pressures for each of the classes have risen and fallen to the present day. If this pattern of rapidly-varying selection for escape from different antibody classes continues, the phenotypic evolution of SARS-CoV-2 may become less predictable in the future than it has been since the appearance of Omicron.

**Figure 6.**
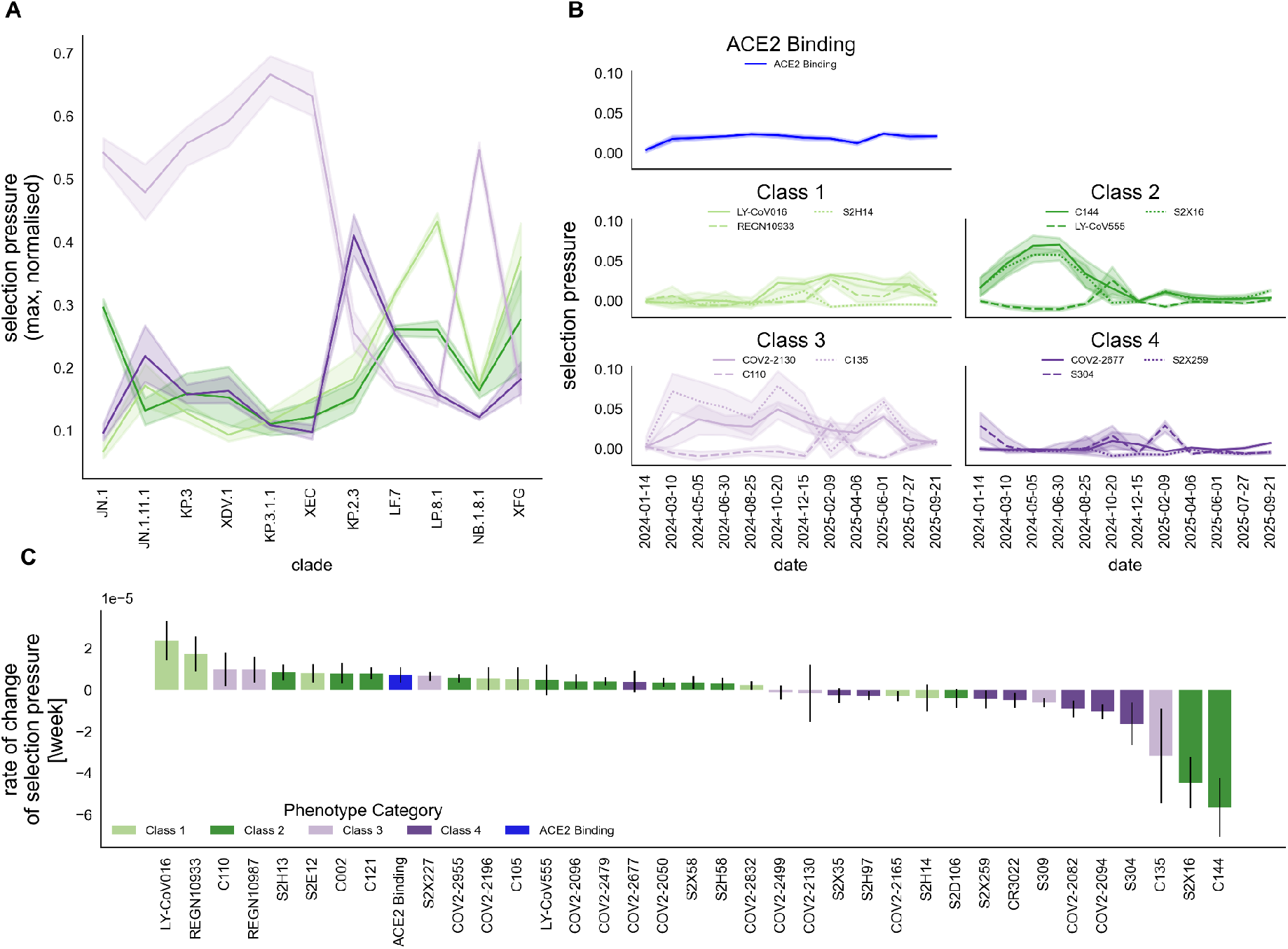
Selection on recent variants circulating in 2024 and 2025. (A) Relative strength of selective pressure for immune escape from different antibody classes, estimated for individual variants ordered by time of emergence. (B) Time trends in selective pressure within the lineage descending from JN.1, estimated for different phenotypes. (C) Rate of change over time in the selective pressure for a panel of phenotypes, estimated across the lineage descending from JN.1.

## Discussion

In this work we have explored for the first time how natural selection has acted on SARS-CoV-2 phenotypes and how these selective pressures have changed in time during the pandemic. This unique study connects to the rich literature on the evolutionary history of SARS-CoV-2, and illustrates the power of combining high-quality data sources – large-scale phylogenetics and multi-phenotype DMS – to understand the drivers of virus evolution. Our work also provides a unique glimpse into the dynamics of a local fitness seascape across different timescales, illustrating how selective pressures changed week after week and year after year during the COVID-19 pandemic.

All but one of the phenotypes in this study are measures of antibody neutralisation, enabling us to build a detailed picture of selection for immune escape. Antibodies belonging to the same Barnes class display similar trends of inferred selection for immune escape, suggesting that the population antigenic landscape induces broadly class-specific selective pressures. We confirm the suggestion^32^ that class 2 is the most important at the start of the pandemic; the same authors also suggested that once the virus has escaped some class 2 antibodies class 1 and 3 will supply the most selective pressure, which is confirmed by our analysis. One of the most interesting changes at the beginning of the pandemic is the shifted immunodominance to class 3 for Beta^33^ which indeed corresponds to the first time during the pandemic that escape from class 3 antibodies appear to be under stronger selection than class 2 in our analysis (Figure 1B). Later on, we observe that natural selection on Omicron is still driven by class 2 escape, but with a stronger role by both class 1 and 3 escape, consistent with computational predictions^34^.

Apart from early Omicron variants (BA.1-BA.5), the most interesting change that we observe in selective pressure has occurred for recent major variants. All these variants (except NB.1.8.1) descend phylogenetically from BA.2.86, which is characterised by strong selection for class 1 escape in addition to class 2. Unsurprisingly, its descendant JN.1 is characterised by massive escape from class 1/2 antibodies, caused by a single amino acid change in the spike protein^35,36^. Selective pressure on JN.1 and KP clades shifted completely to class 3 antibody escape, with a complete and unprecedented drop off in all class 2 escape fitness gradients. This shift was predicted on an immunological basis by Paciello et al.^35^.

The antibody escape component of the seascape is particularly dynamic. SARS-CoV-2 immune escape appears to follow a Red Queen pattern^37^: immune pressure leads to emergence of a new immune escape variant, the human immune system adapts to the different antigenic profile of this variant, leading to the emergence of new variants with different antigenic profile, and so on. This process is clearly visible when comparing the temporal dynamics of selective pressures with the sequence of evolutionary changes of SARS-CoV-2.

Selective pressures and population composition can be used to forecast short-term evolution within SARS-CoV-2 clades. Most importantly, selective pressures can predict longer-term evolution, In fact, we demonstrate a clear link between time-varying selective pressures (i.e. changes in the fitness seascape for a particular phenotype) and SARS-CoV-2 evolution (i.e. changes in that phenotype that are fixed in major variants). This holds for several individual molecular phenotypes and supports the idea that once major variants emerge from persistent infections, their success is driven by similar selective pressure as minor variants generated during acute infections. This link provides predictive power to our approach, enabling probabilistic prediction of phenotypes in future major variants. Although these predictions are inherently noisy, we are able to effectively estimate the probability of substantial phenotypic shifts in newly emerging variants.

Our initial hypothesis was that the selective pressure on ACE2 binding would have weakened after the emergence of Omicron, while pressure for immune escape would have significantly increased. In fact, natural selection for immune escape is already strong for early variants but becomes stronger after BA.1 as expected, albeit with complex dynamics across different antibody classes. This evolutionary transition to escape-dominated selection mirrors the observed epidemiological transition to purely variant-driven dynamics ^38^. Interestingly, there is no evidence of decrease in selective pressure for ACE2 binding. ACE2 binding is critical for cell entry, therefore some selective pressure should be expected. A possible explanation for the relatively constant pressure for increased ACE2 binding is compensatory evolution^39^: mutations conferring antibody escape are pleiotropic and tend to reduce ACE2 binding, therefore moving this phenotype far away from the optimum and increasing the fitness gradient, which favour the emergence of compensatory mutations restoring the initial binding affinity. However, the observed increase in time of selective pressure within clades is not fully consistent with this explanation.

Other works have also attempted to quantify the contribution of different phenotypes to fitness. Ma et al. used multivariate regression to determine the contribution of immune escape and ACE2 binding to mutation fitness, finding that ACE2 binding was more often the better predictor^27^. However, they only explored a limited number of major clades, leaving time trends unclear, and considered ‘immune escape’ as merely one aggregated phenotype from deep mutational escape data across many different antibodies. Maher et al. explicitly explored time trends in the predictive power of phenotypic, (as well as epidemiological and other) features on the spread of mutations^17^. They found that ACE2 binding’s predictive power rose and fell, peaking in the summer of 2020, while antibody escape (here derived from changes in simulated binding energies across a range of antibodies) peaked later in early 2021. However, their temporal analysis was completed before the emergence of Omicron. Dadonaite et al. found that for XBB.1.5 sub-clades, sera escape DMS data was the better predictor of clade growth, compared to ACE2 binding and cell entry^16^. Most recently, Haddox et al. have directly explored the different patterns of evolution for escape from hundreds of antibodies, identifying common temporal patterns in the changing levels of escape^40^. Haddox et al. link some of these escape patterns to the spread of individual mutations, and for a small number of these mutations explore whether the mutation spread patterns are indicative of changing selection (i.e. changing fitness) or clonal interference (selection/fitness does not change). Meanwhile, in our study we focus on a much smaller range of antibodies with available deep mutational scanning data with high coverage across mutations. This allows us to obtain estimates of the selective pressure (fitness gradient) for escape for the individual antibodies, and how it changes. We also show that viral adaptation within a clade does not necessarily proceed only through those mutations that later define new clades: in fact, the selection signal can be observed also on non-clade-defining mutations only (Supplementary Figure S9).

Our work has a few limitations. First, mutations are often pleiotropic and can cause escape from multiple antibodies. Just because a fitness gradient is high does not mean there is a selective pressure to escape that specific antibody; rather, it could be a pleiotropic consequence of a selective pressure to escape a different one. A similar issue applies for evolutionary changes: just because the virus escaped a particular antibody does not mean it was under selection, since it could be a by-product of selection on some other phenotype and resultant changes. However, by classifying phenotypes based on biological characteristics such as Barnes epitope classes, we remove some of this ambiguity and examine clear signals in the changing fitness landscape.

Novel strains will also diverge from the original strains in their sequence, and therefore explore different parts of the fitness landscape. The effect of previous evolutionary changes on the structure of the local landscape is known as epistasis and is an important driver of changes in evolutionary directions. A form of epistasis that we already discussed is diminishing returns^28^: once several mutations that enhance a phenotype have been fixed, there is less advantage in enhancing it further. This form of epistasis leads to decreasing fitness gradients while fitness increases. However, epistatic interactions can be far more complex ^41^: for example, the huge immune escape of Omicron was only tolerable because of its increased ACE2 receptor binding affinity coming from N501Y and Q498R ^42^. We are able to account for epistasis in our phenotypic data on ACE2 binding as there exists data for several different genetic backgrounds, but such multi-background data is not generally available in antibody binding escape studies. Hence, temporal changes in the seascapes we obtain could be partly driven by epistatic interactions.

In addition, we are able to study only a limited number of phenotypes, therefore we cannot fully resolve the pleiotropies. More importantly, we cannot detect selective pressure on non-measured phenotypes. For example, we expect that the late evolution of SARS-CoV-2 after BA.2.86 will be driven by class 3 escape, and possibly even class 4; however, since we do not have DMS studies for antibodies that are relevant for these late variants, we cannot fully detect the corresponding selection and therefore we underestimate the selection strength. This effect partly explains the late decrease in selection for immune escape in Figure 1B. However, the biases due to pleiotropy also partly compensate for this effect, since patterns of selection can be revealed by other antibodies of the same class included in the DMS studies, even if these antibodies are not directly selected.

During future major epidemics of emerging viruses or new strains, we expect that DMS studies and large-scale sequencing will be rapidly deployed. A key strength of our work is the ability to infer the selective pressures on the virus – i.e. the local phenotypic fitness landscape – from these measurements. Since natural selection is the driver of evolution, this approach will provide valuable information to understand future evolutionary changes before they happen. For example, based on recent selective pressures (Figure 6), we can expect a short-term shift of the circulating viral population towards increased escape from class 1 antibodies, and possibly class 2 antibodies (if the next variants will descend from XFG) or class 3 (if from NB.1.8.1). More generally, this work establishes a framework for real-time inference of selective pressures from sequence and phenotypic data, providing a forward-looking view of how viral pathogens adapt to shifting immune landscapes.

## Acknowledgments

We are thankful to Jesse Bloom, Richard Neher and Dalan Bailey for useful discussions and insights. CH was supported by funding from the Biotechnology and Biological Sciences Research Council (UKRI-BBSRC) [grant number BB/T008784/1]. We acknowledge support from the European REA, Marie Skłodowska-Curie Actions, grant agreement no. 101131463 (SIMBAD). This work was funded by UK Research and Innovation (UKRI) under the UK government’s Horizon Europe funding guarantee [grant number EP/Y037375/1].

