## Supplementary Information for "Natural selection driven by escape from shifting antibody classes shapes SARS-CoV-2 evolution"

Charlie Hamilton et al.

#### S1 Supplementary Materials and Methods

##### S1.1 Phylogenetic fitness estimator

The analysis in this paper is founded on comparing mutations' $\Delta f$ (fitness effect) and $\Delta \phi_i$ (phenotypic effect on a phenotype $i$). To estimate the fitness of individual mutations, we use a modified version of the Bloom and Neher phylogenetic fitness estimator `SARS2-mut-fitness` [1]. Their method estimates the expected number of counts $n_{\text{expected}}$ of the occurrence of a mutation (as if it were under no selection, which should hold for four-fold degenerate sites) and compares that with the actual number of mutations counts inferred from the phylogenetic tree $n_{\text{actual}}$. A mutation's fitness effect is then given by

$$\Delta f = \log\left(\frac{n_{\text{actual}} + P}{n_{\text{expected}} + P}\right), \tag{S1}$$

where $P$ is a constant pseudocount equal to 0.5.

###### S1.1.1 Phylogenetic data

Bloom and Neher's fitness estimator method is applied to mutation-annotated phylogenetic trees from [2]. The particular phylogenetic tree we use includes all public sequences (from "GenBank, COG-UK, and the China National Center for Bioinformation") up to 1st September 2025 (available at http://hgdownload.soe.ucsc.edu/goldenPath/wuhCor1/UShER_SARS-CoV-2/2025/09/01). Subsets of this tree are then used for the following applications.

###### S1.1.2 Inter-clade analysis

In our analysis of how the fitness landscape changes between clades we calculate the fitness effect of all possible mutations, in each of the Nextstrain clades [3] we consider. In Bloom and Neher's original method, the expected and actual count of a mutation are both calculated on a clade-by-clade basis, and hence $\Delta f$ is already defined relative to a specific Nextstrain clade. The clades that feature in our analysis are those with a sufficient number of sequences, and hence sufficient mutation counts to draw accurate results. We set the cutoff to 2500 sequences per clade; the cutoff is set at this level to allow the most recent clades e.g. 25B (NB.1.8.1) to be included in the analysis. For clades that have more sequences than this cutoff, we uniformly downsample the sequences in that clade's phylogenetic tree to this number (with 10 independent downsamples to determine the uncertainty resulting from this downsampling).

###### S1.1.3 Intra-clade analysis, explicit time variation

For the analysis of the variation of the fitness landscape within one clade, over time, an additional step is necessary to calculate the fitness effect $\Delta f$ for each mutation at a particular time period. First, for a particular clade, we determine the number of time periods of a particular duration (say 1 week) in which the number of sequences available exceeds a certain cutoff value of $N_{\text{cutoff}}$ sequences. Both the time period and the sequence cutoff are hand-picked to give approximately 10 different time periods for a clade. A particular time period $i$ has start time $t_{\text{start},i}$ and end time $t_{\text{end},i}$. The earliest sequence in the clade has associated time $t_0$.

For each time period, we take two subsections of the clade's phylogenetic tree, spanned by a subset of sequences starting at $t_0$. The first subsection (A, 'the bigger') includes sequences with times $t_0 \leq t < t_{\text{end},i}$. The sequences in this subsection are uniformly downsampled by the ratio

$$r = \frac{N_{\text{cutoff}}}{N[t_{\text{start},i} \leq t < t_{\text{end},i}]},$$

where $N[t_{\text{start},i} \leq t < t_{\text{end},i}]$ is the number of sequences in the time range $t_{\text{start},i} \leq t < t_{\text{end},i}$. This ensures that each of these A subsections has approximately $N_{\text{cutoff}}$ sequences in the period of interest $t_{\text{start},i} \leq t < t_{\text{end},i}$. We independently downsample 10 times.

For each of these subsections A, we also take a further subsection (B, 'the smaller'). This subtree B is spanned by the same sequences in the range $t_0 \leq t < t_{\text{start},i}$ that are also included in its A counterpart. This construction means there should be approximately $N_{\text{cutoff}}$ sequences in subsection A that are not in B, and that have associated times within the time period of interest.

We can then determine the expected and actual counts of each mutation in both section A and B. We calculate

$$n_{\text{actual}} = n_{\text{actual,A}} - n_{\text{actual,B}}$$

and similarly for $n_{\text{expected}}$. The fitness effects $\Delta f$ are then calculated for each mutation according to equation (S1). The way we have constructed these subsections A and B means these mutation counts occur in the time period of interest ($t_{\text{start},i} \leq t < t_{\text{end},i}$).

#### S1.2 Phenotypic data

The phenotypic effect values $\Delta\phi_i$ come from deep mutational scanning (DMS) data. We have chosen datasets with full coverage of mutations in the Receptor Binding Domain (RBD) of the Spike protein to maintain consistency throughout. Our main source of antibody escape DMS data is a compiled resource available at `https://github.com/jbloomlab/SARS2_RBD_Ab_escape_maps/blob/main/processed_data/escape_data.csv` [4]. This dataset contains escape data for mutations across many different antibodies. We limit our analysis to the data which comes from Jesse Bloom or Tyler Starr labs, as these have much higher mutation coverage than those from Yunlong Cao (since we are interested in determining aggregate quantities relating to the whole landscape, not just specific mutations, it is essential to have as high a coverage as possible). We are left with escape data for 36 different antibodies, each of which is measured against a Wuhan-Hu-1 background, and which are also labelled according to their Barnes epitope class. Data on ACE2 binding is from Taylor and Starr [5], and includes a range of backgrounds.

#### S1.3 Estimating phenotypic contribution to fitness

As described in the Main Text, the phenotypic fitness landscape admits a linear description around a set of phenotypes that do not correspond to a peak in the landscape. Considering a time-dependent landscape $f(\phi, t)$, the fitness values can be approximated around a reference set of "wild-type" phenotypes $\phi^{wt}$ by a Taylor expansion,

$$f(\phi, t) = f(\phi^{wt}, t) + \sum_i a_i(\phi^{wt}, t)\left(\phi_i - \phi_i^{wt}\right) + \text{higher order terms} \tag{S2}$$

or, in short,

$$\Delta f(\phi^{wt}, t) = \sum_i a_i(\phi^{wt}, t)\Delta\phi_i + \text{higher order terms.} \tag{S3}$$

The fitness effects vary both explicitly with time $t$, and with the genetic background (which note may also vary with time). We can take a genetic background $g$ as reference, instead of a set of phenotypic values, i.e. consider $\phi^{wt} = \phi(g)$ and fitness effects $\Delta f = \Delta f(g, t)$. For the phenotypes we consider (i.e. antibody escape, ACE2 binding), the phenotypic effect of a mutation $\Delta\phi_i$ is a biophysical quantity and should be independent of time, but may depend on the genetic background, $\Delta\phi_i = \Delta\phi_i(g)$. Therefore, we have that

$$\Delta f(g, t) = \sum_i a_i(g, t)\Delta\phi_i(g) + \text{higher order terms.} \tag{S4}$$

As a consequence, if we consider a mutation $\mu$ with phenotype effects $\Delta\phi_{i,\mu(g)} = \phi_i(g_\mu) - \phi_i(g)$ and fitness effect $\Delta f_\mu(g) = f(g_\mu) - f(g)$, the above equations result in

$$\Delta f_\mu(g, t) = \sum_i a_i(g, t)\Delta\phi_{i,\mu} + \text{higher order terms} \tag{S5}$$

It follows that the strength of the selection pressure for a particular phenotype $i$ is also given by the relevant coefficient $a_i$ in the Taylor expansion of the fitness effect of a particular mutation.

We wish to determine the time variation of the phenotypic contributions to fitness, where such contributions are given by the expansion coefficients $a_i(g, t)$. For practical reasons, instead of the independent variation with $t$ and $g$ we will consider the variation with the major clade/variant $c$ to investigate long-term evolutionary trends since the emergence of SARS-CoV-2. The sequence of major clades has a time order, so can be treated as a proxy for time $t$. Choosing to explore the variation with the major clade introduces another effect because the genetic background will vary $g = g(c)$, but we do not expect this contribution to dominate. We also explore the case of explicit time variation within single clades, and hence where the genetic background does not vary much, $g \approx g_0$.

While phenotypic effects can change with the genetic background (due to epistasis), the mutational effect on many antibody escape phenotypes are measured against just one background, giving a constant $\Delta\phi_i(g) = \Delta\phi_i$. For ACE2 binding there is data available for multiple backgrounds, so we can include this dependence to some extent. For a particular genetic background $g$, we use the phenotypic effects whose background is the most recent direct ancestor of $g$.

For both variation with major clade and explicitly with time, we wish to determine the expansion coefficients $a_i(t)$ at different times. We do this by extracting the partial regression coefficients. We perform a linear Ridge regression with standardised dependent and independent variables (standardised to mean zero and unit variance), with regularization parameter $\alpha = 10^3$. Regularization is necessary because there is generally substantial correlation between the effects of mutations on different phenotypes (mutations are pleiotropic). Example Ridge trace plots in Figure S6 shows how the values of the regression coefficients vary for changing regularization $\alpha$. We observe that for values greater than $\alpha \approx 10^3$ the coefficients are more stable – especially their relative order.

#### S1.4 Combining selective pressures

We combine selective pressures from antibodies, both altogether and according to their Barnes binding class, in order to understand their total effect. In the Main Text, this is accomplished simply be taking the maximum selection across all the particular antibody escape pressures belonging to the particular category. This is a simple method that is independent of the number of phenotypes.

Another possible method is to aggregate as follows. We have a set of fitness gradients $a_i$ for phenotypes belonging to some category, with associated phenotypic effects $\Delta\phi_i(m)$ for a mutation $m$. We wish to make an equality

$$a_{tot}\Delta\phi_{tot}(m) = \sum_i a_i \Delta\phi_i(m) \tag{S6}$$

Now, from our standardization of the phenotypic variables entering into the regression we have that

$$\sum_n \Delta\phi_i(n) = 0, \tag{S7}$$

and

$$\mathrm{Var}_n(\Delta\phi_i(n)) = 1. \tag{S8}$$

It follows that these conditions should also to apply to our constructed $\phi_{tot}$. One such expression is

$$\Delta\phi_{tot}(m) = \frac{\sum_i a_i \Delta\phi_i(m)}{\mathrm{Var}_n(\sum_i a_i \Delta\phi_i(n))}. \tag{S9}$$

This quantity clearly has zero mean and unity variance (over all mutations). It then follows from equation S6 that

$$a_{tot} = \mathrm{Var}_n\left(\sum_i a_i \Delta\phi_i(n)\right). \tag{S10}$$

In Figure S2, we plot the results of aggregating selection pressures in this manner, compared to taking the maximum as in Main Text Figure 1B/C.

#### S1.5 Model for antibody escape selection dynamics

In Figure 4 (Main Text) we show a model of antibody selection dynamics. This is a multi-strain SIR model (see e.g. [6]). In particular, we consider four different strains. All the strains have the same infectivity $\beta$, and recovery rate $\gamma$. They are distinguished by exactly which susceptible compartments they can infect:

1. The original 'wildtype' strain ($w$) that only infects naive individuals ($S$)
2. A 'neutral' strain ($n$) that can also infect recovereds from the wildtype ($S + R_w$)
3. A 'partial escape' strain ($e$), that can infect a fraction $k$ of recovereds from neutral strain, recovereds from wildtype, and naive individuals ($S + R_w + kR_n$)
4. A 'complete escape' strain ($c$) that can reinfect all the recovereds from previous strains, plus naive individuals ($S + R_w + R_n$)

For any infected individual, the infecting virus may mutate to a new strain with a rate $\mu$. We assume that mutations between strains occur one step at a time, and only in a specific order of increasing escape, i.e. $w \to n \to e \to c$

(equivalently we can label these numerically from 1 to 4). The SIR-like dynamical equations are the following,

$$\frac{dS}{dt} = -\beta S\left(I_w + I_n + I_e + I_c\right) \tag{S11}$$
$$\frac{dI_w}{dt} = \beta S I_w - \gamma I_w - \mu I_w \tag{S12}$$
$$\frac{dI_n}{dt} = \beta(S + R_w)I_n - \gamma I_n + \mu I_w - \mu I_n \tag{S13}$$
$$\frac{dI_e}{dt} = \beta(S + R_w + kR_n)I_e - \gamma I_e + \mu I_n - \mu I_e \tag{S14}$$
$$\frac{dI_c}{dt} = \beta(S + R_w + R_n)I_c - \gamma I_c + \mu I_e \tag{S15}$$
$$\frac{dR_w}{dt} = \gamma I_w - \beta R_w(I_n + I_e + I_c) \tag{S16}$$
$$\frac{dR_n}{dt} = \gamma I_n - \beta R_n(kI_e + I_c) \tag{S17}$$
$$\frac{dR_e}{dt} = \gamma(I_e + I_c) \tag{S18}$$

The model as described allow us to define and calculate concepts to describe the selection dynamics. The fitness of a particular strain is described by its effective reproductive number, which depends on its particular available susceptible fraction. That is,

$$f_w = \frac{\beta}{\gamma}S \tag{S19}$$
$$f_n = \frac{\beta}{\gamma}(S + R_w) \tag{S20}$$
$$f_e = \frac{\beta}{\gamma}(S + R_w + kR_n) \tag{S21}$$
$$f_c = \frac{\beta}{\gamma}(S + R_w + R_n) \tag{S22}$$
$$\tag{S23}$$

The existence of varying fitness for different genotypes yields a fitness landscape. In turn, this fitness landscape depends on the antigenic landscape of the population, i.e. the amount of susceptibles to different strains $(S, R_w, R_n)$. Define fitness difference between consecutive strains such that

$$\Delta f_i = f_{i+1} - f_i \tag{S24}$$

At time $t$, the prevailing fitness gradient in the infected population (i.e. how much of a benefit there is to evolving a new mutation) depends on the relative abundances of the different strains and their fitness differences. It is given by

$$\Delta f(t) = \frac{1}{\sum_j I_j}\sum_j I_j \Delta f_j \tag{S25}$$

i.e., a weighted average of the fitness differences weighted by the strain prevalences.

We are interested in the phase of the epidemic that starts with a major wave caused by the neutral strain. For this reason, we define the prevailing escape (that is, escape from the *neutral* strain that is shaping the antibody landscape at the point of interest) prevalence as a weighted average of those escape fractions,

$$E = \frac{1}{\sum_j I_j}(kI_e + I_c) \tag{S26}$$

Similarly, we define the population immunity (to the *neutral* strain) as the sum of the relevant recovered fractions,

$$R_{\text{pop}} = R_n + R_e + R_c. \tag{S27}$$

These recovered fractions act as a source of new susceptibles for either the partial escape strain, the complete escape strain, or a future strain that can also escape the latter two.

#### S1.6 Predictive model for antibody escape

We develop a model for predicting change in antibody escape in a clade from the historic selection pressure acting on that particular antibody. That is, we wish to predict the phenotypic change $\Delta\phi_i(g)$ where $i$ denotes the phenotype in question and $g$ is the new, clade-defining genotype. This phenotypic change is defined with respect to some 'reference' phenotype,

$$\Delta\phi_i(g) = \phi_i(g) - \phi_i^{ref}(g) \tag{S28}$$

where $\phi_i(g)$ is the value of the phenotype for the clade-defining genotype, given by the sum of the phenotypic effects of its defining mutations,

$$\phi_i(g) = \sum_{m \in g} \Delta\phi_i(m) \tag{S29}$$

and the reference value of the phenotype $\phi_i^{ref}(g)$ is taken to be the mean average of the values of the phenotypes of the clades that emerged less than one year before the clade of interest,

$$\phi_i^{ref}(g) = \frac{\sum_{g' \text{ emerged } \leq \text{ 1y before } g} \phi_i(g')}{\sum_{g' \text{ emerged } \leq \text{ 1y before } g}}. \tag{S30}$$

Similarly, we define the historic selection pressure for a phenotype $i$ acting on a clade $g$, $a_i^{hist}(g)$ as the mean average of the selection pressures that acted on clades that emerged less than one year previously,

$$s_i^{hist}(g) = \frac{\sum_{g' \text{ emerged } \leq \text{ 1y before } g} a_i(g')}{\sum_{g' \text{ emerged } \leq \text{ 1y before } g}}, \tag{S31}$$

and $a_i(g')$ is determined by the Ridge regression of mutation fitness effects against phenotypic effects as described in Section S1.3.

Our predictive model is then trained and evaluated on all pairs $(a_i^{hist}(g), \Delta\phi_i(g))$. We consider a two-stage model with both a logistic part and a linear part. This two-stage model is motivated by the dynamics of evolution by natural selection: the logistic part accounts for the fact that small selection pressures may not be translated into meaningful evolutionary change, due to stochastic effects. If a meaningful evolutionary change does arise, its value should be (to first order) a linear function of the selection. Schematically, then, we assume a functional relationship,

$$\Delta\phi_i(g) = \text{logistic}(a_i^{hist}(g)) \cdot \text{linear}(a_i^{hist}(g)), \tag{S32}$$

i.e. the linear part only comes into play if the logistic part equals unity, otherwise the value of the whole function is zero.

Here 'meaningful/substantial' denotes any phenotypic change $\Delta\phi_i(g)$ whose magnitude is greater than a particular threshold, $\epsilon$ (i.e. $-\epsilon \leq \phi_i(g) \leq \epsilon$). The variation of the proportion of phenotypic changes which are substantial with the historic selection pressure, for different values of the threshold $\epsilon$, is shown in Figure S10. In our model we set $\epsilon = 0.1$, as this value yields proportions of phenotypic changes which are substantial that cover a sensible range (beginning at $\sim$0.2 for low selection values and rising to $\sim$0.8 for high selection).

The structure of our assumed functional relationship means the two parts of the regression can be accomplished separately. We binarise the values of phenotypic change (according to the threshold criteria above) and perform a logistic regression on these binarised values. For these pairs $(a_i^{hist}(g), \Delta\phi_i(g))$ where $-\epsilon \leq \phi_i(g) \leq \epsilon$), we then perform ordinary least squares linear regression of the phenotypic change against the selection.

In the Main Text, we show the performance of the two-stage regression model both when it is trained and evaluated on the entire dataset, and using a conventional 80/20 train/test split. We summarise the accuracy of these two instances of the model below:

| Training source | Test source | Baseline log loss | Logistic part loss (Cross-entropy) | Linear part $R^2$ |
|---|---|---|---|---|
| All | All | 0.664 | 0.592 | 0.0197 |
| Training set (80%) | Test set (20%) | 0.656 | 0.558 | 0.0164 |

**Table S1:** Performance of the predictive model for antibody escape from historical selection pressure.

In both cases, the loss of the logistic part of the regression (cross-entropy loss) is smaller than the baseline, showing improved ability to predict whether a phenotypic change will be substantial from the selection pressure. The $R^2$ of the linear part of the regression is small, though positive. However this is because a lot of the general positive relationship between the selection pressure and the phenotypic change is instead being captured by the logistic part. If one simply does OLS regression of the phenotypic change (without filter) and the selection pressure, $R^2 = 0.0797$.

### S2 Supplementary Results

#### S2.1 Clade-by-clade analysis

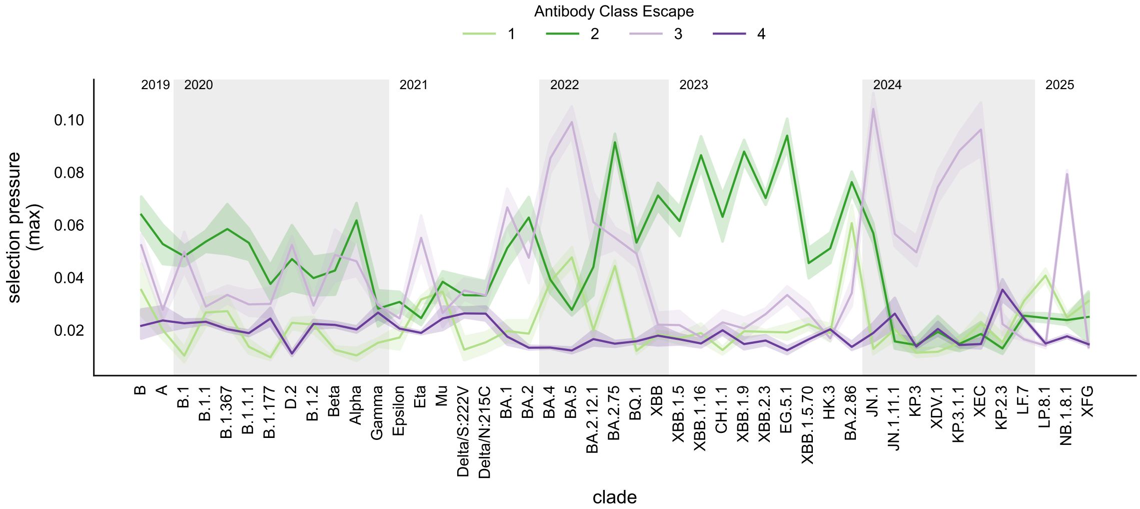


**Figure S1:** Maximum selection pressures for escape for antibodies within each Barnes class, shown for each Nextstrain clade considered in our main analysis. This figure is identical to Figure 2C in the Main Text, except that here the selection pressures are not normalised.

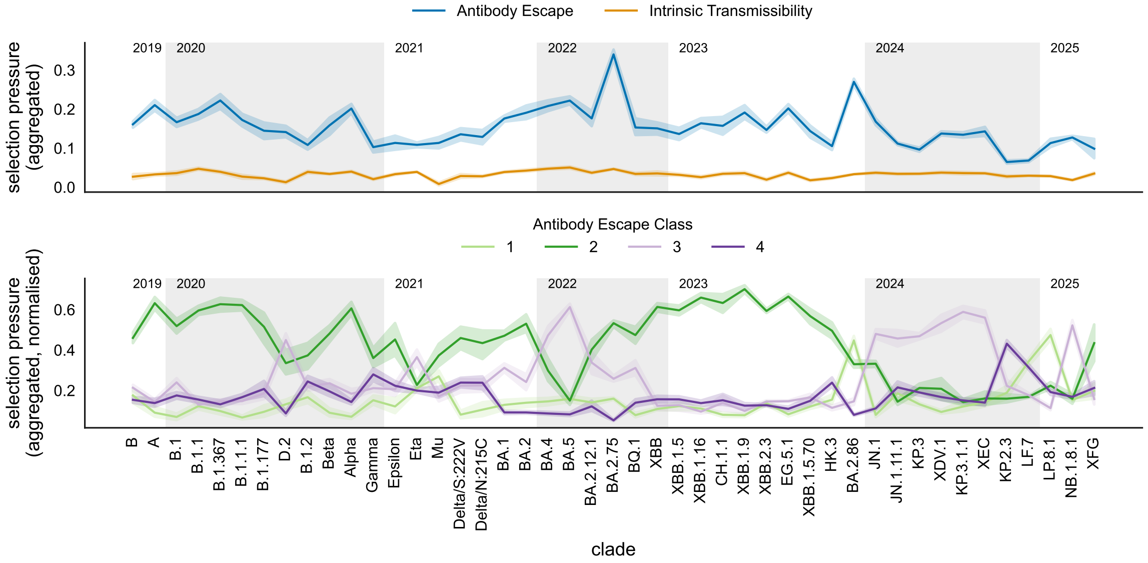


**Figure S2:** Aggregated selection pressures according to equation (S10), for all clades. This figure is identical to Main Text Figure 1C, appart from aggregating in this manner instead of taking the maximum selection pressure across phenotypes in the class. (Top) phenotypes are split according to being either antibody escape / intrinsic transmissibility. (Bottom) antibody escape phenotypes are split according to their Barnes class.

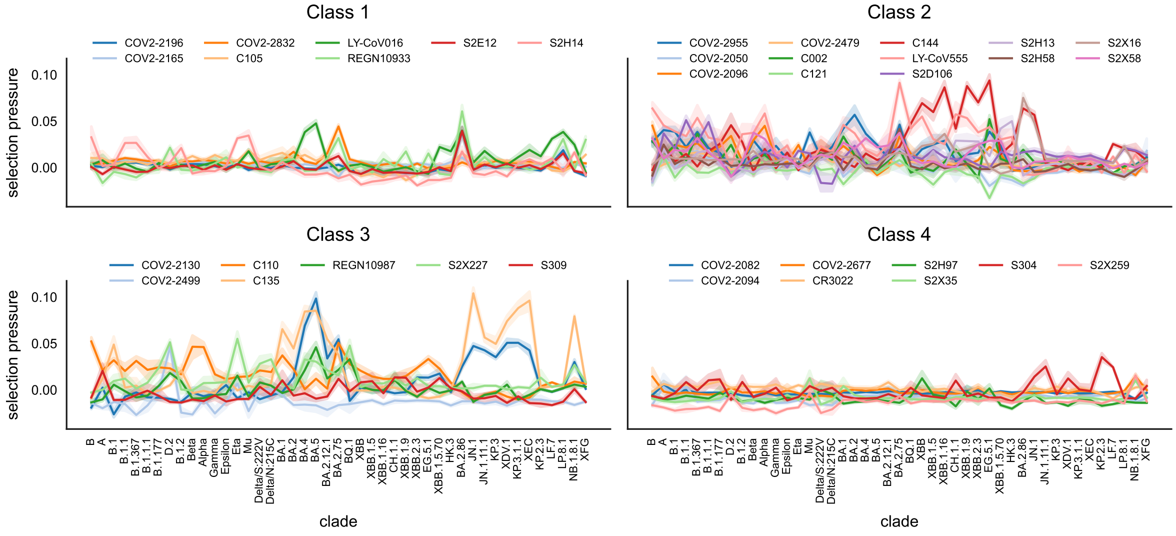


**Figure S3:** Selection pressure for all antibody escape phenotypes, grouped by their Barnes class, as measured in all Nextstrain clades considered.

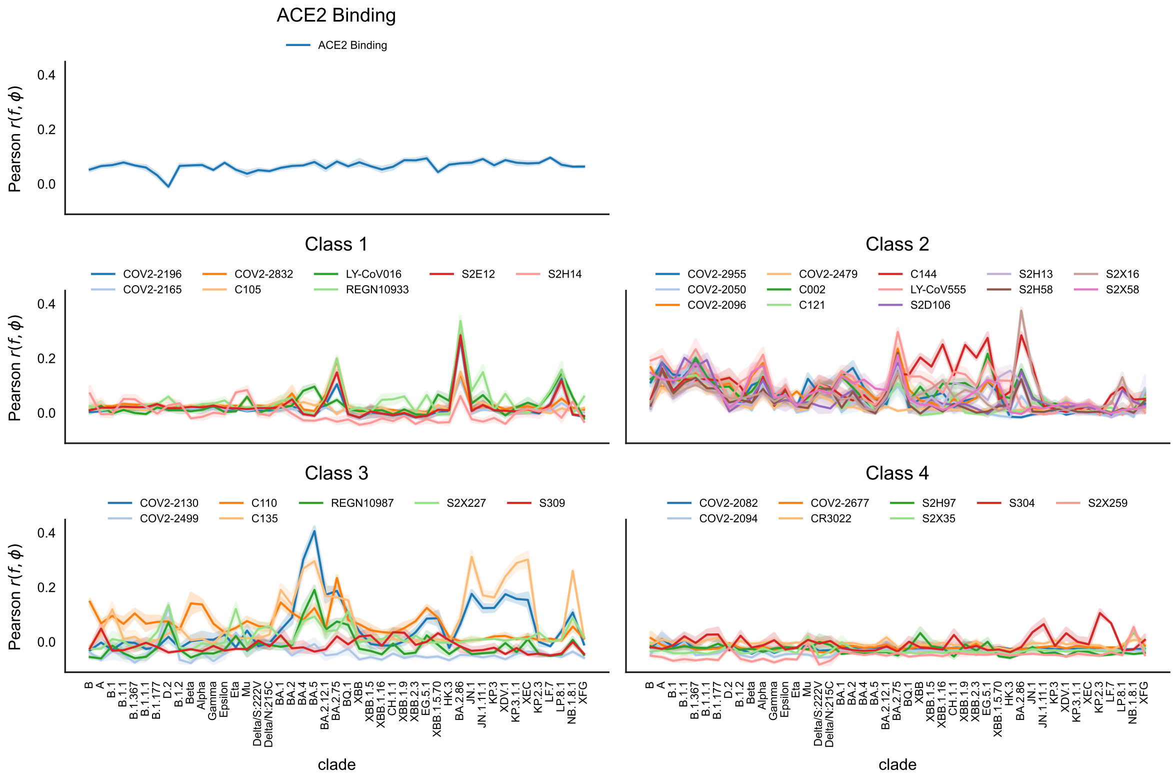


**Figure S4:** Pearson correlation coefficient between fitness and phenotype mutation effects, shown for each Nextstrain clade we consider, and for each of the main phenotypes considered in our analysis. Antibody escape phenotypes are grouped according to their Barnes class.

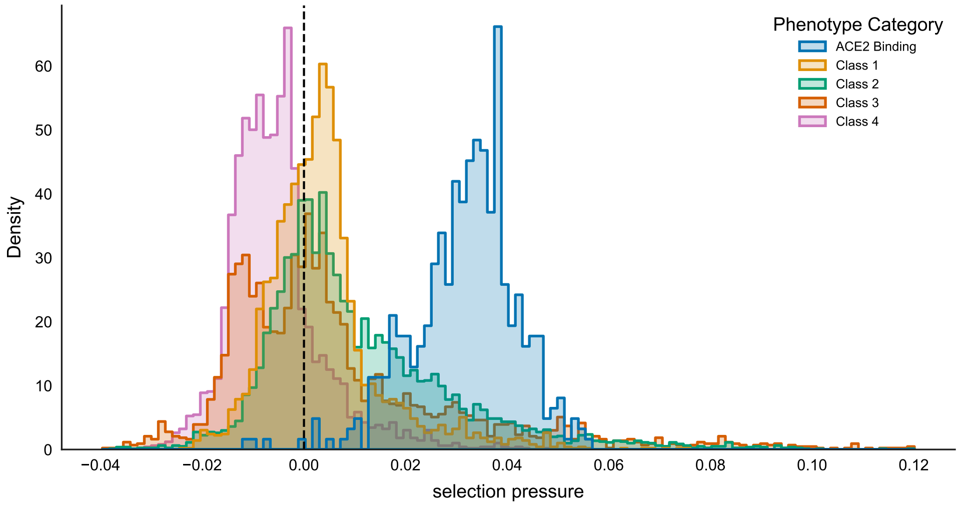


**Figure S5:** The distribution of values of the measured selection pressure (across all phenotypes, clades and repeat downsamples) as determined by the Ridge regression. Phenotypes are classified as being either ACE2 binding, or escape from one of the four Barnes antibody classes.

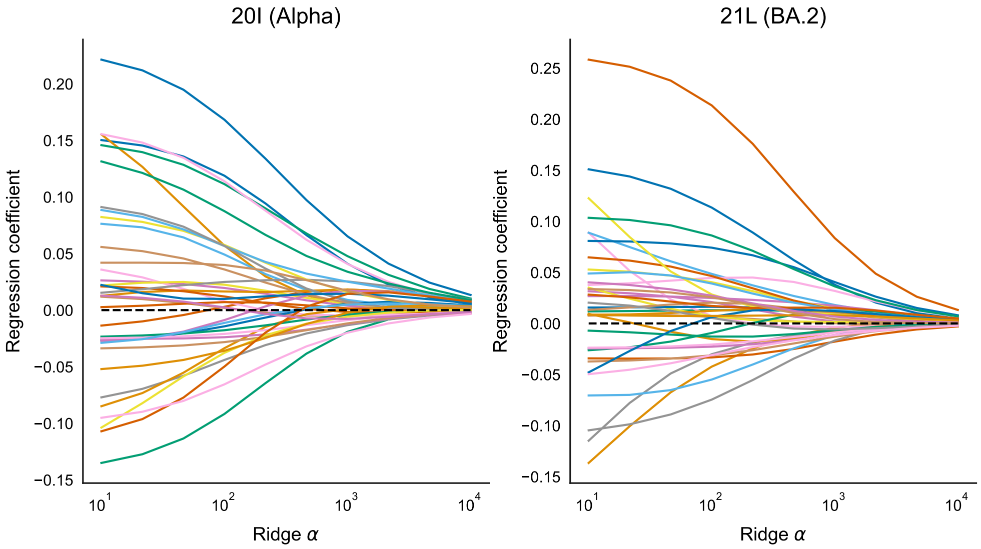


**Figure S6:** Example Ridge trace plots, showing the variation of the partial regression coefficients against each phenotype with the regularization coefficient $\alpha$. Here, the fitness values come from the Alpha/Nextstrain 20I clade (Left) and Omicron BA.2/Nextstrain 21L (Right), each for first of the 10 random downsamples. It can be seen in both examples that for values greater than $\alpha \approx 10^3$, the regression coefficients are more stable.

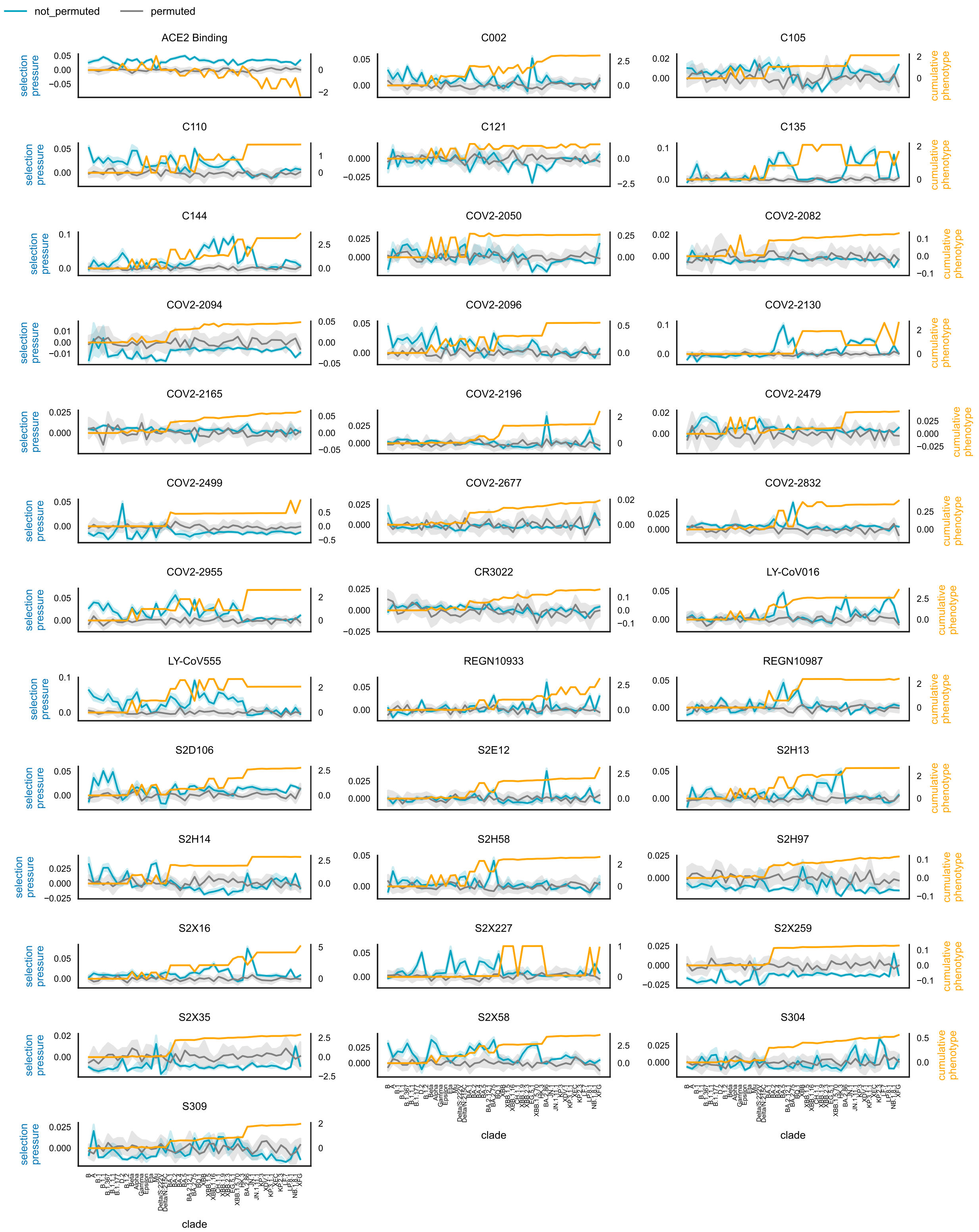


**Figure S7:** Comparison of selection pressures for a phenotype measured within a clade (blue) with the realised phenotype of the clade-defining sequence (orange). This realised phenotype is given by the sum of the phenotypic effects of the defining mutations of the clade founder. Also shown is the selection pressure calculated for the fitness values randomly permuted across different mutations (grey) – this gives an indication of the magnitude of the selection pressure to exceed to be truly indicative of selection.

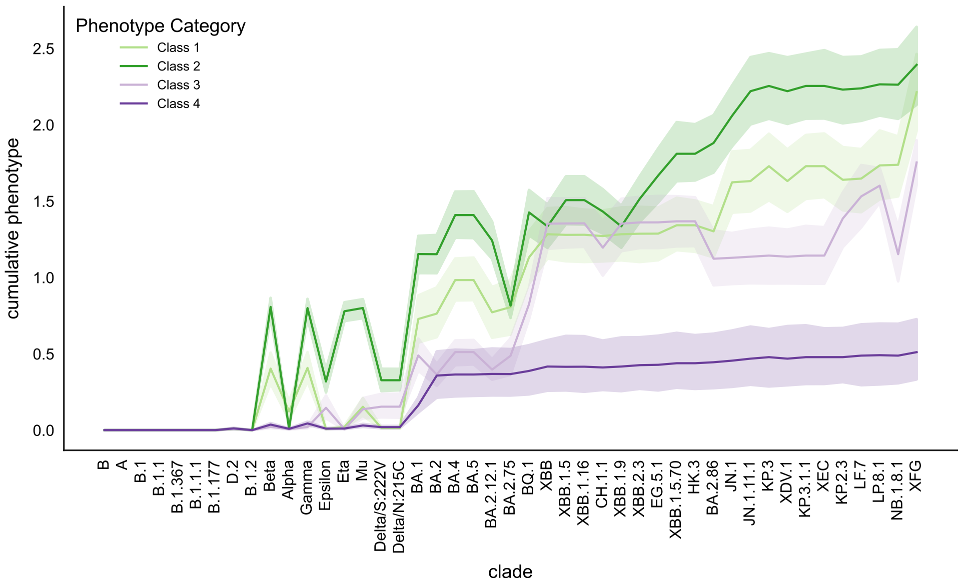


**Figure S8:** Changes in realised escape from each of the four Barnes antibody classes. Shown is the mean escape across antibodies within that class, where for each antibody the value of the phenotype is given by the sum of the phenotypic effects of the clade's defining mutations.

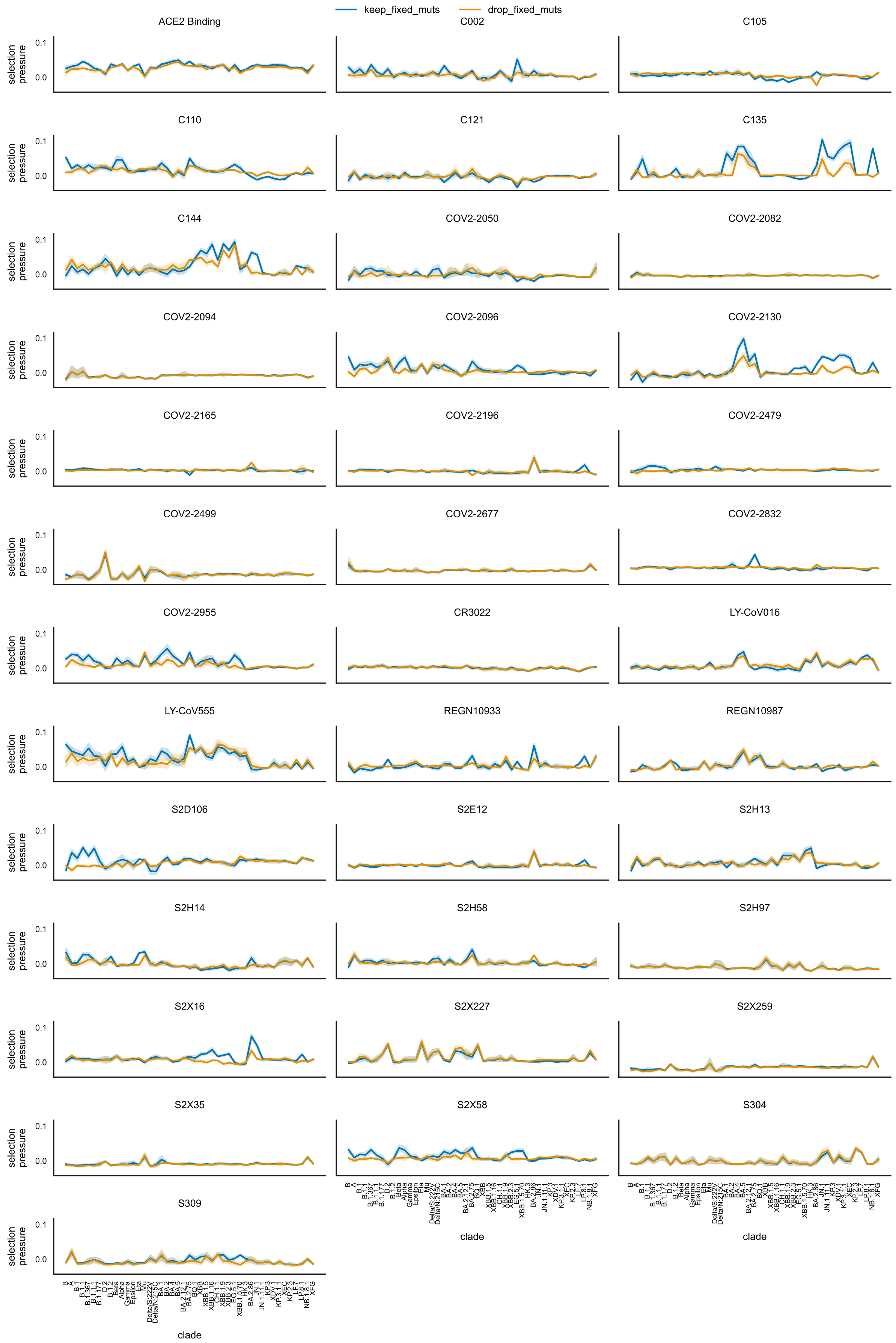


**Figure S9:** The effect of including 'fixed' mutations (mutations that eventually become fixed in some Nextstrain clade, not just the one being considered) on the selection pressure. Shown is both the selection pressure with fixed mutations (blue) and without (orange), for all phenotypes across all clades.

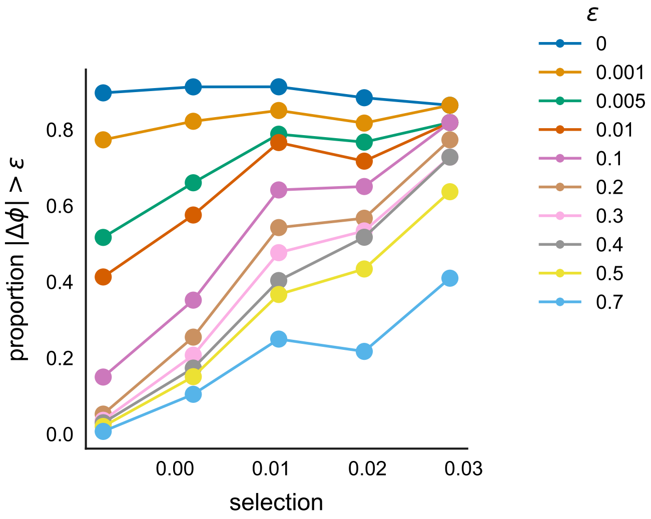


**Figure S10:** The varying proportion of changes in antibody escape $\Delta\phi$ whose magnitude is greater than a particular threshold $\epsilon$, as historic selection pressure for escape from the particular antibody changes. These proportions are computed across all antibodies and clades in our main analysis.

#### S2.2 Time-explicit analysis

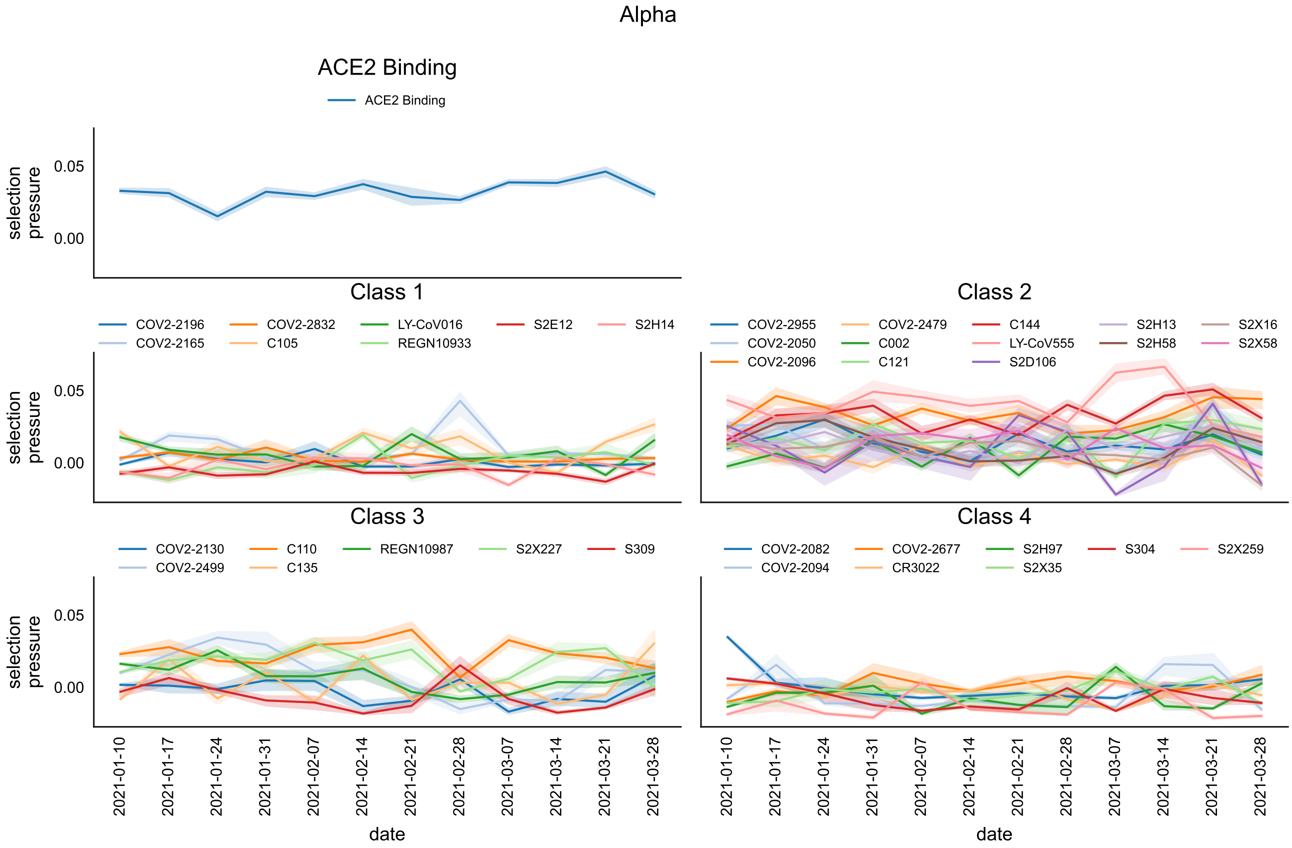


**Figure S11:** Variation of selection pressures for each phenotype with time within the Alpha variant (Nextstrain clade 20I) in England. Phenotypes are arranged according to their class (either 'ACE2 Binding' or antibody escape from one of the four Barnes antibody classes).

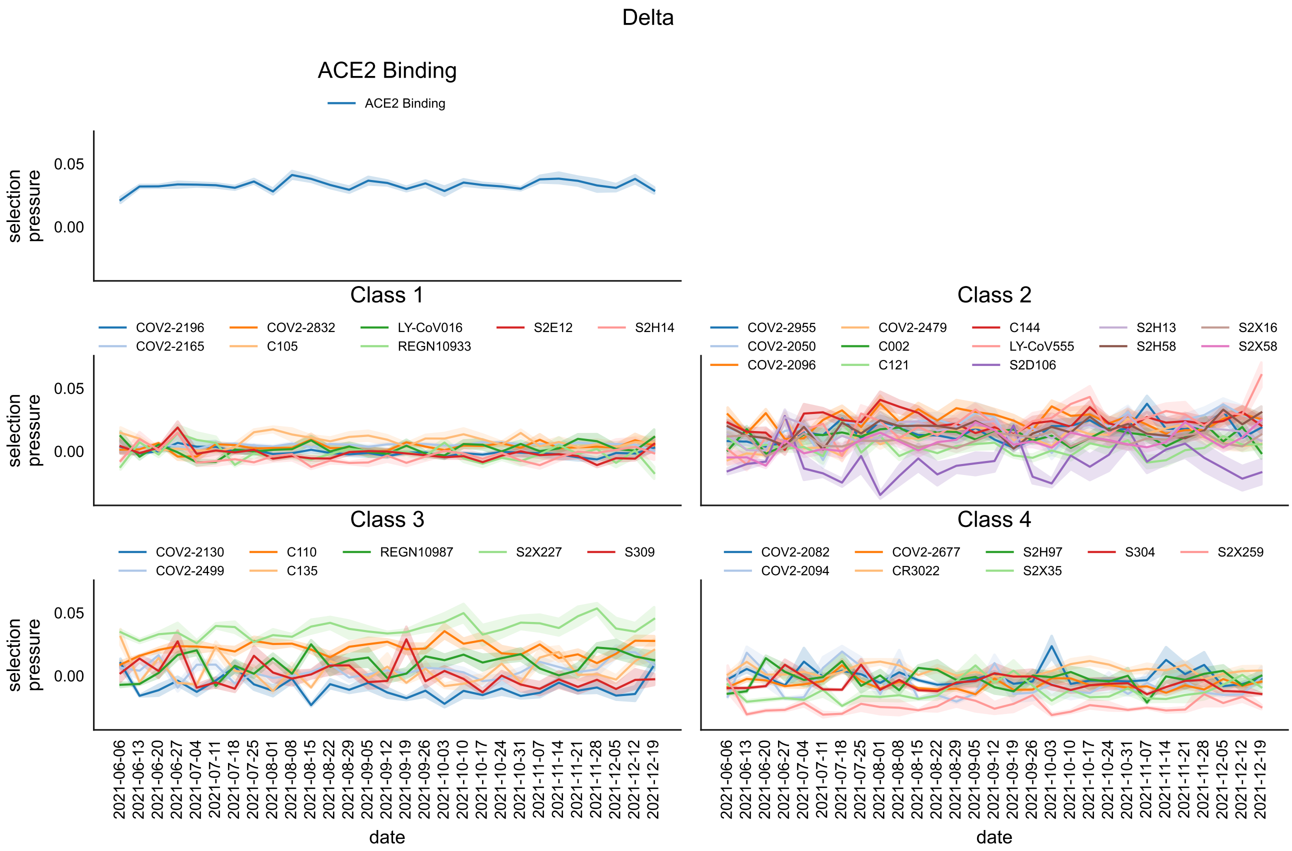


**Figure S12:** As above for the Delta variant (Nextstrain clade 21J).

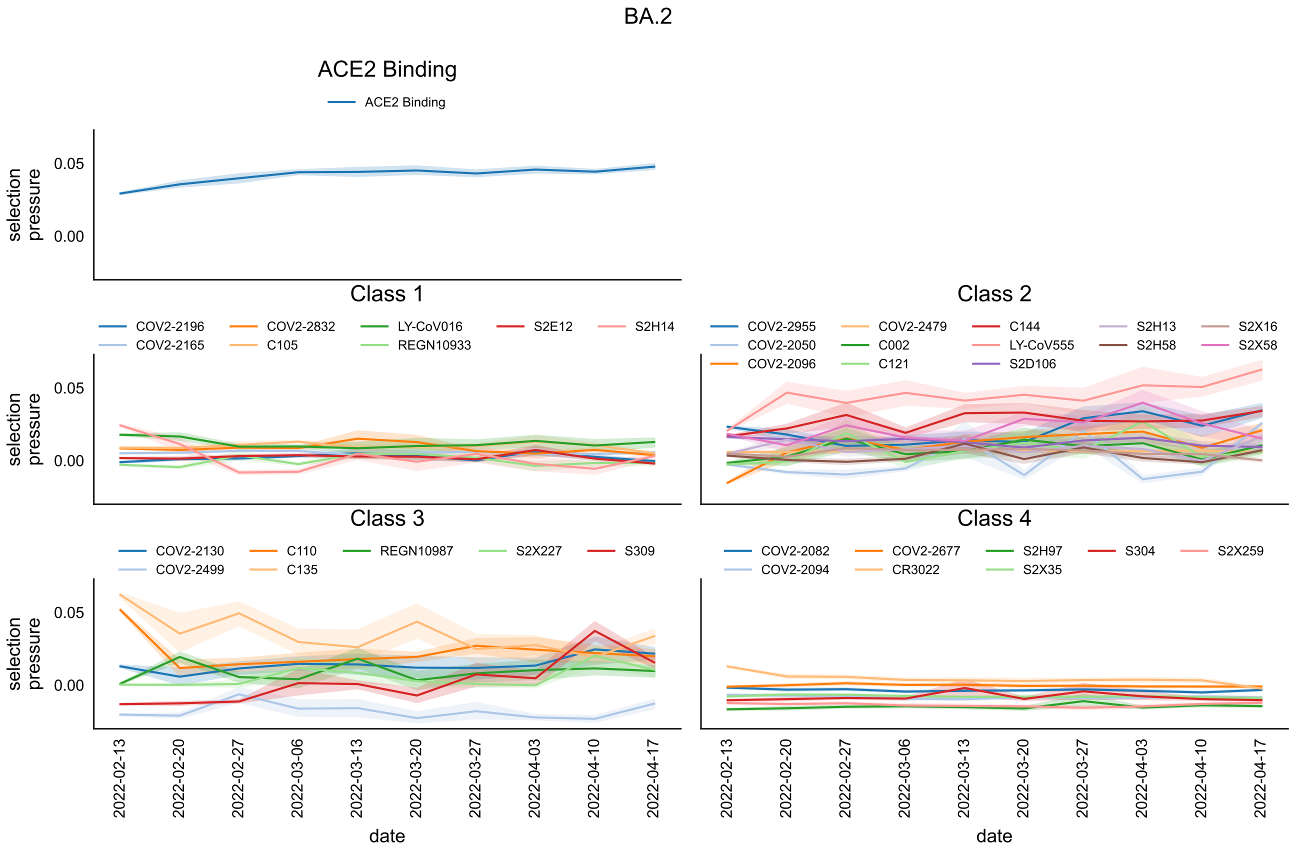


**Figure S13:** As above for the BA.2 variant (Nextstrain clade 21L).

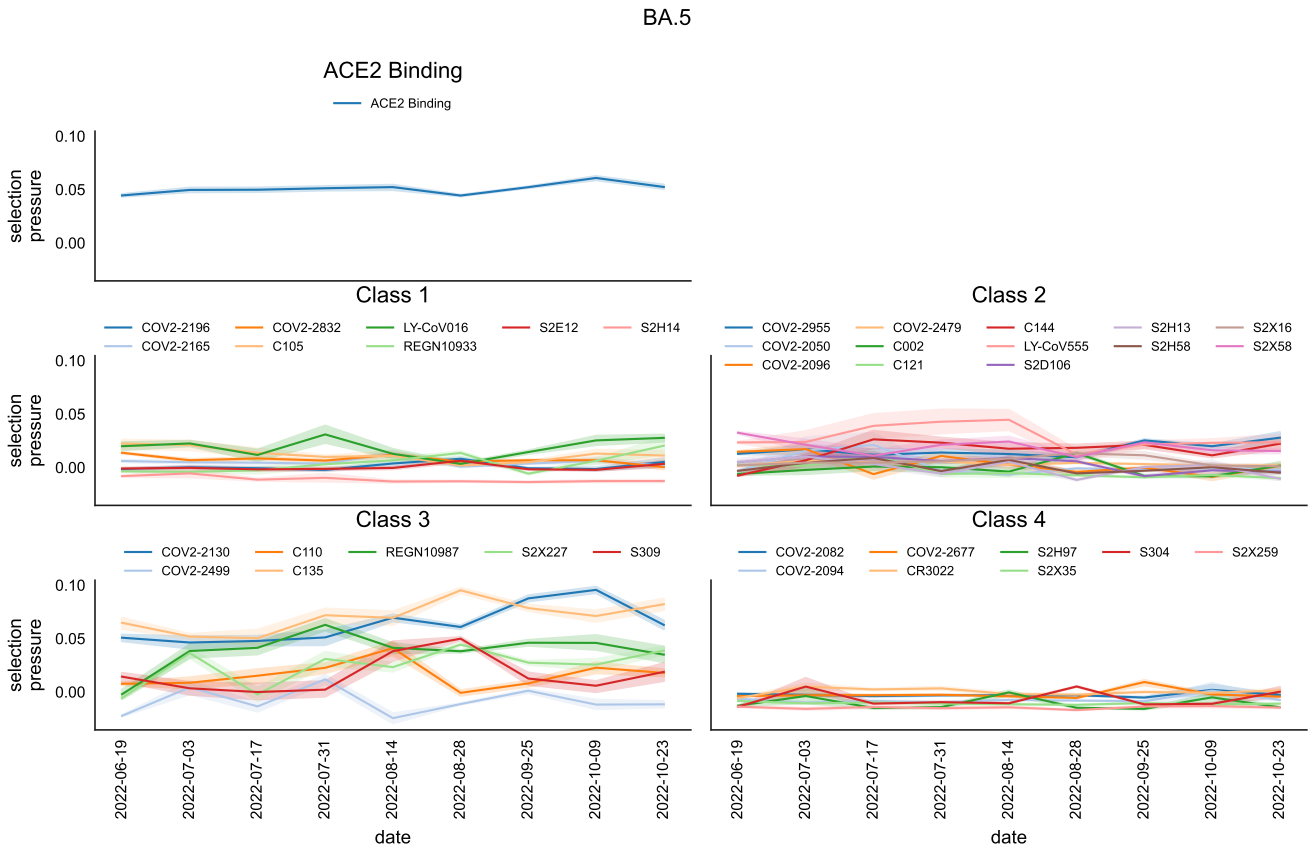


**Figure S14:** As above for the BA.5 variant (Nextstrain clade 22B).

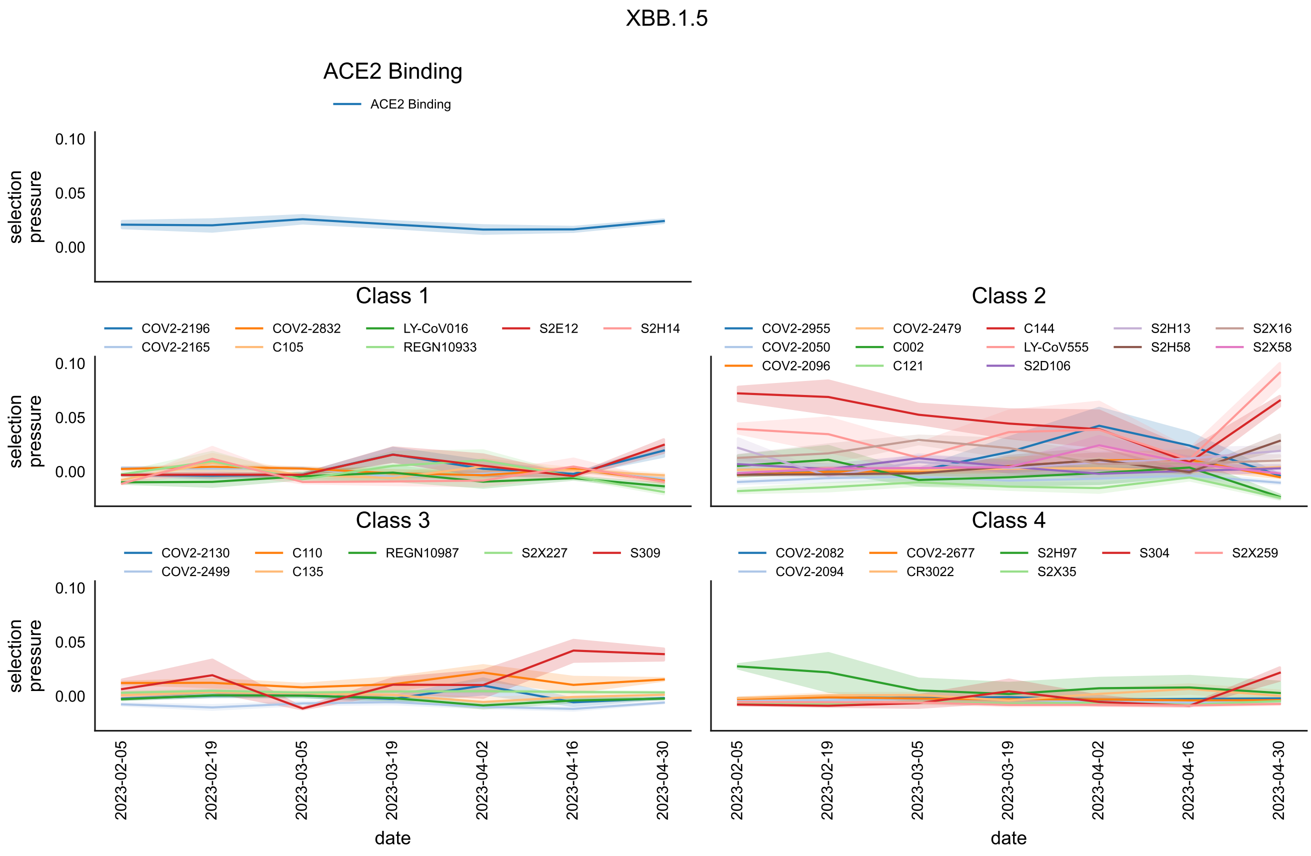


**Figure S15:** As above for the XBB.1.5 (Nextstrain clade 23A).

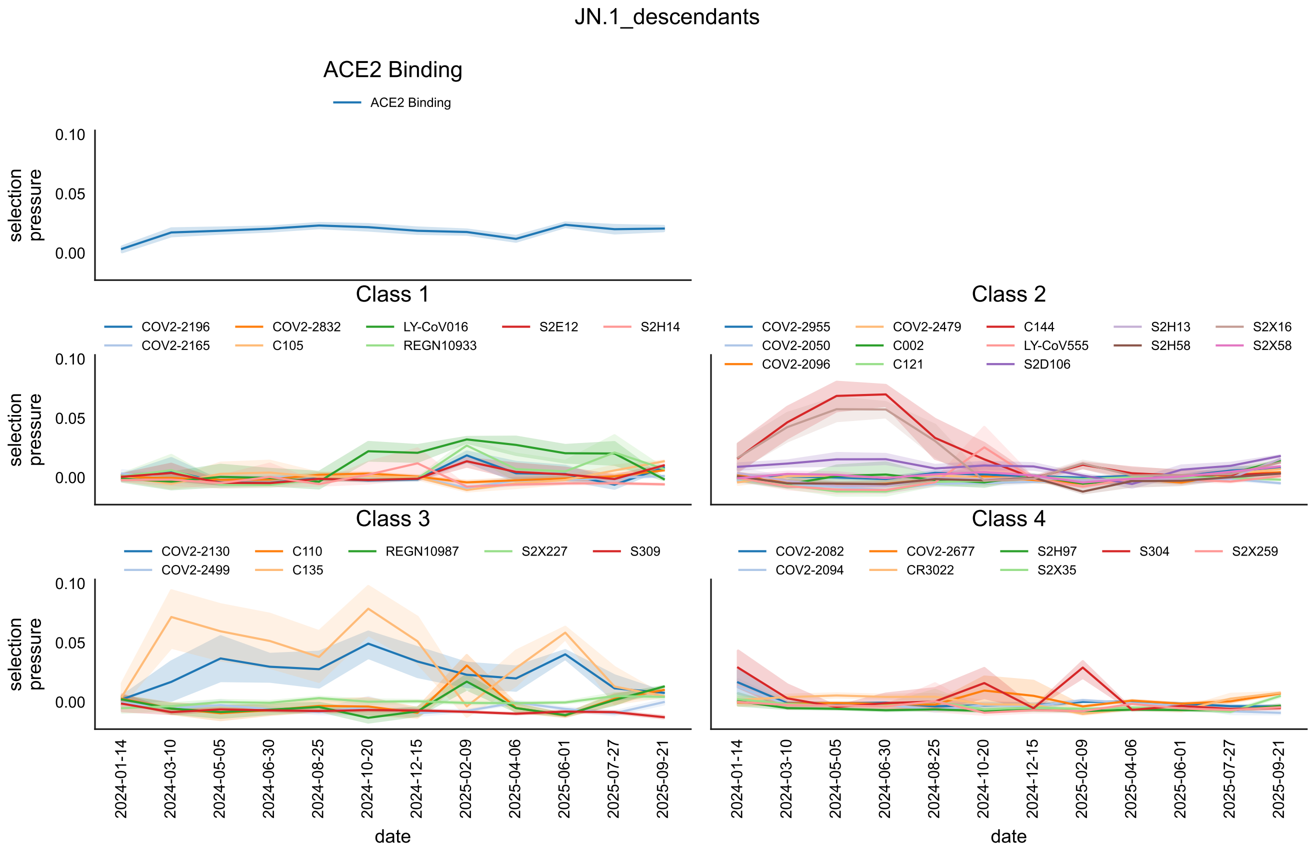


**Figure S16:** As above for the JN.1 variant (Nextstrain clade 24A) and its direct descendants (Nextstrain clades 24B, 24C, 24E, 24F, 24G, 24H, 24I, 25A, 25C).

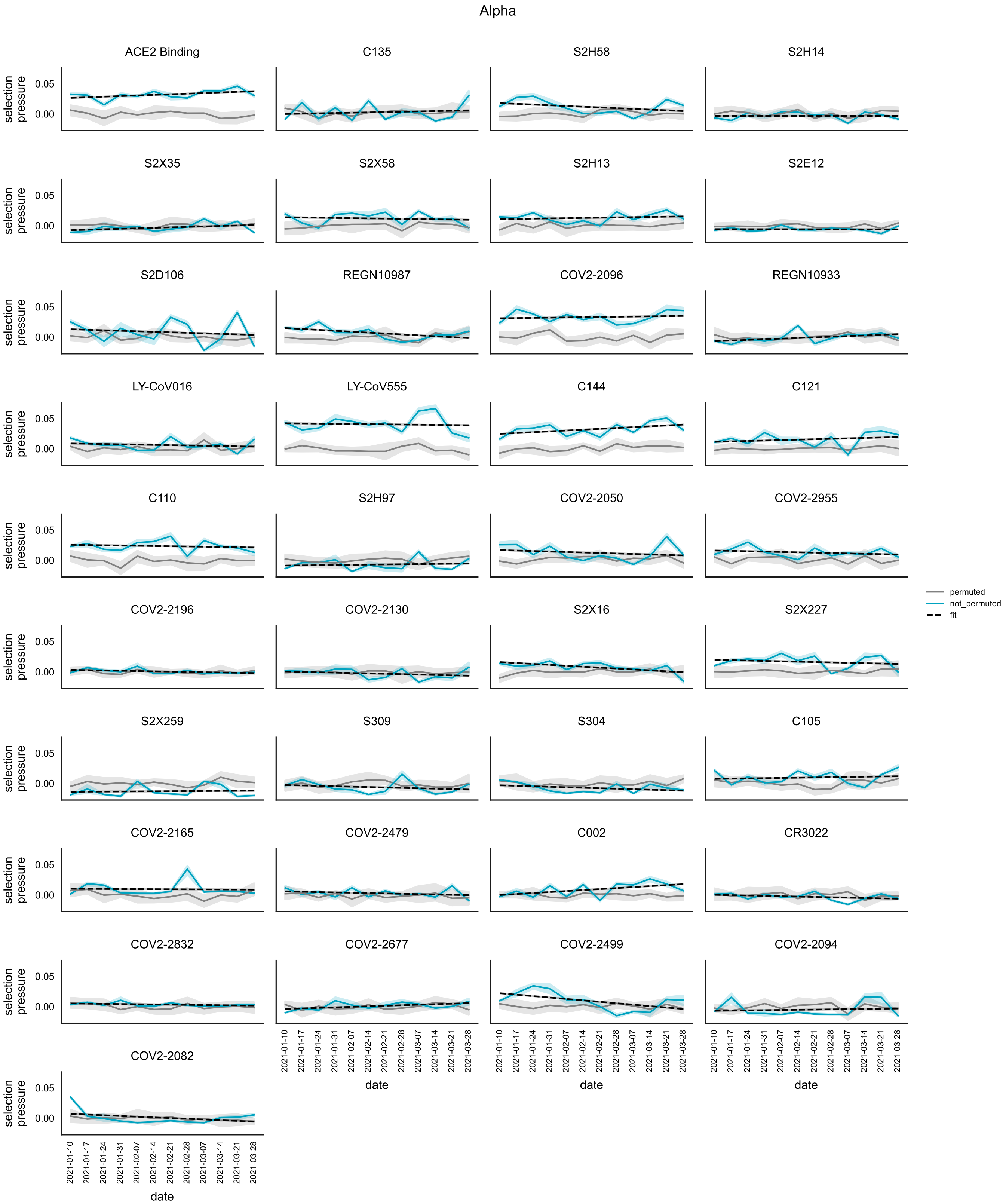


**Figure S17:** Variation of selection pressures (blue) for each phenotype with time within the Alpha variant in England. Also shown is a straight line fit from which the slope/rate of change is extracted (black, dashed) and the selection pressure re-calculated with fitness values have been randomly permuted across mutations (grey).

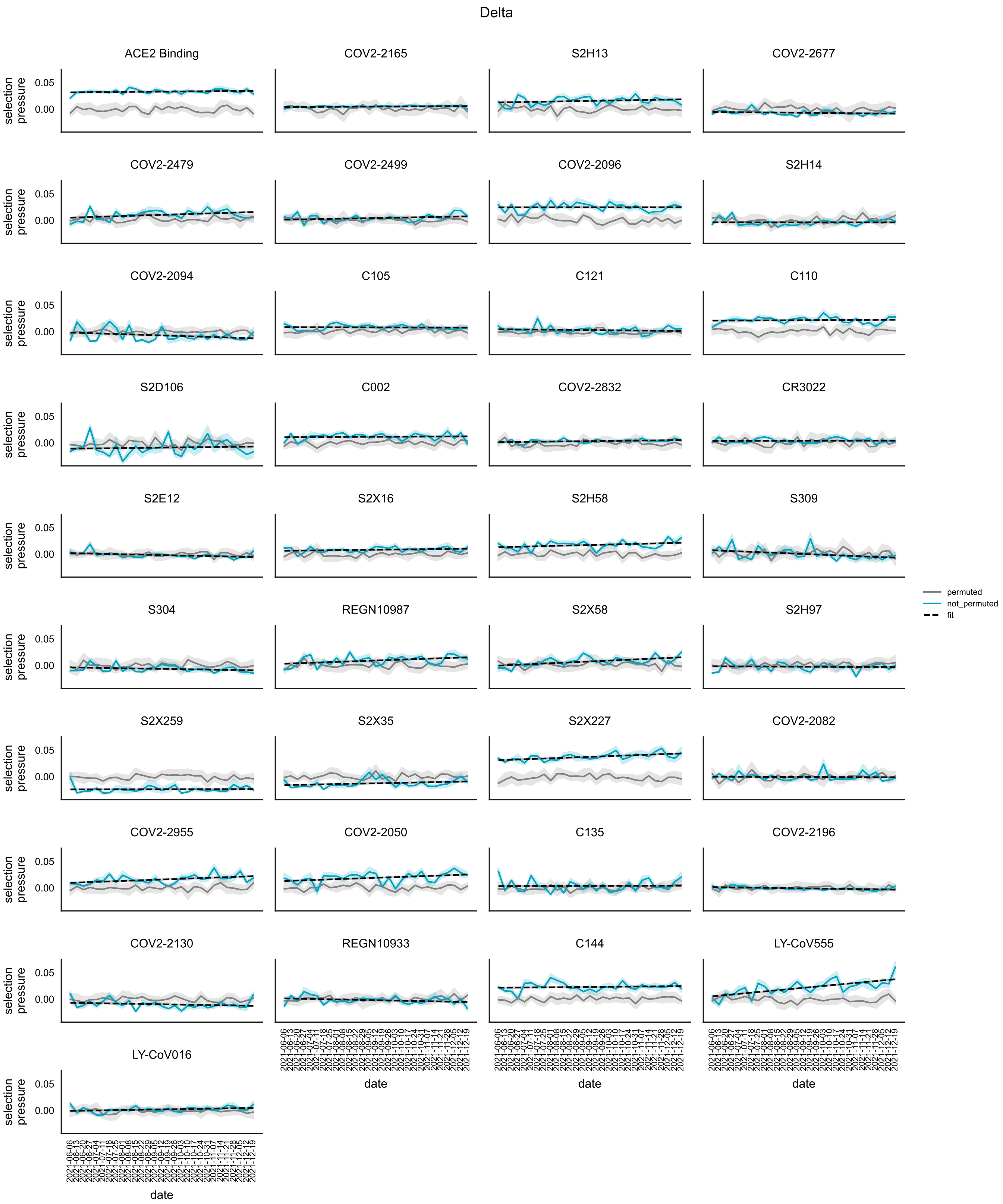


**Figure S18:** As above for the Delta variant (21J).

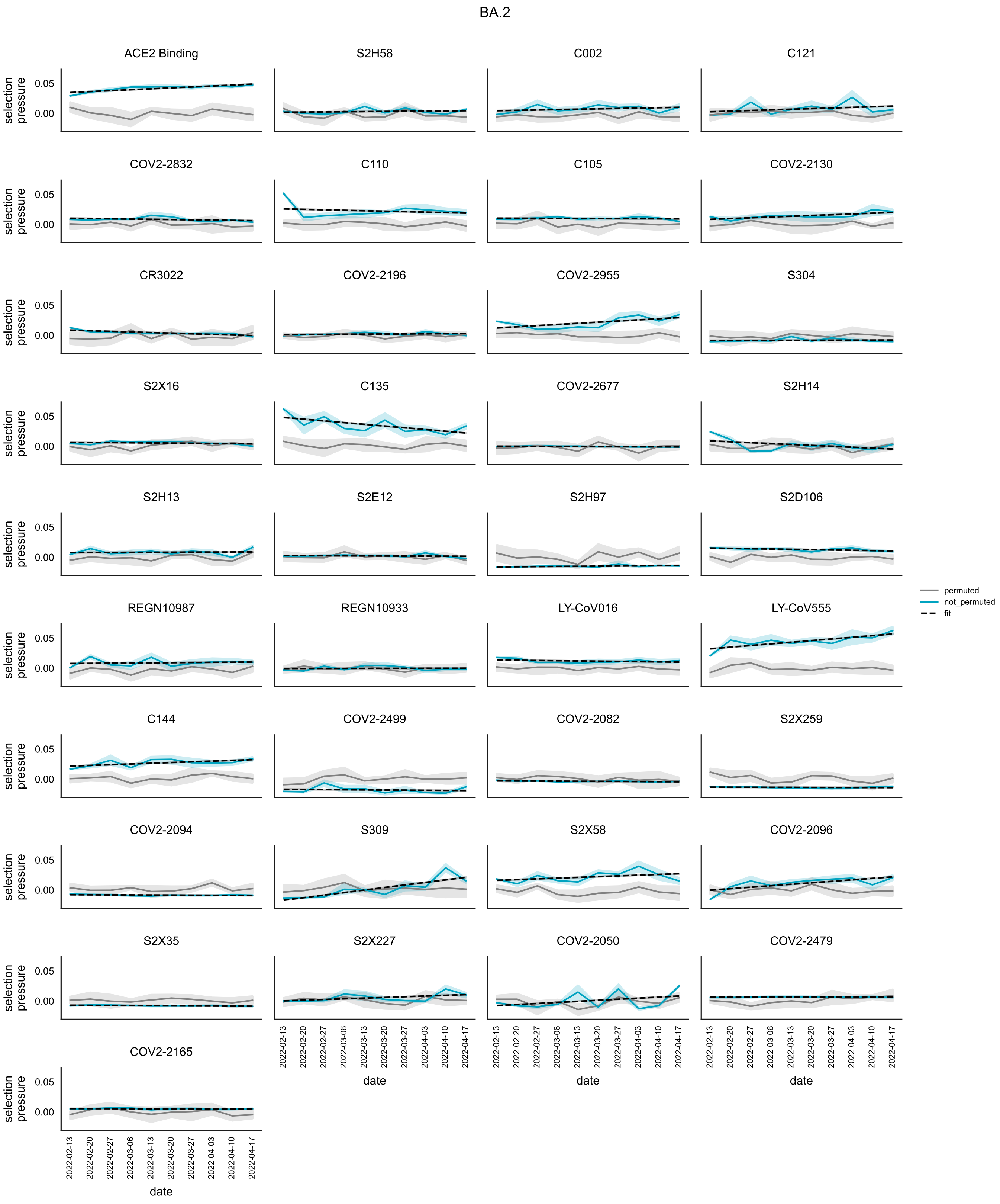


**Figure S19:** As above for the BA.2 variant.

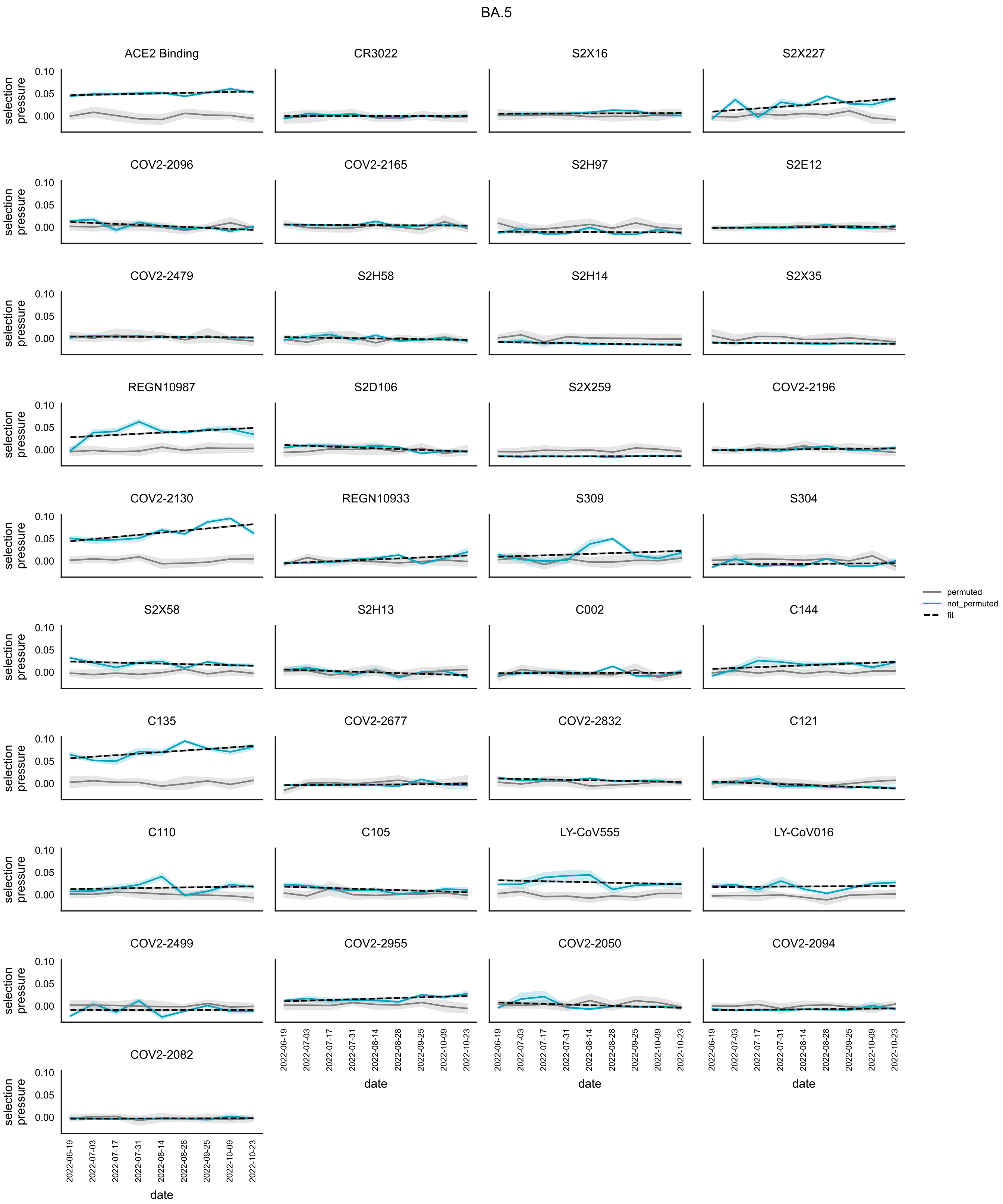


**Figure S20:** As above for the BA.5 variant

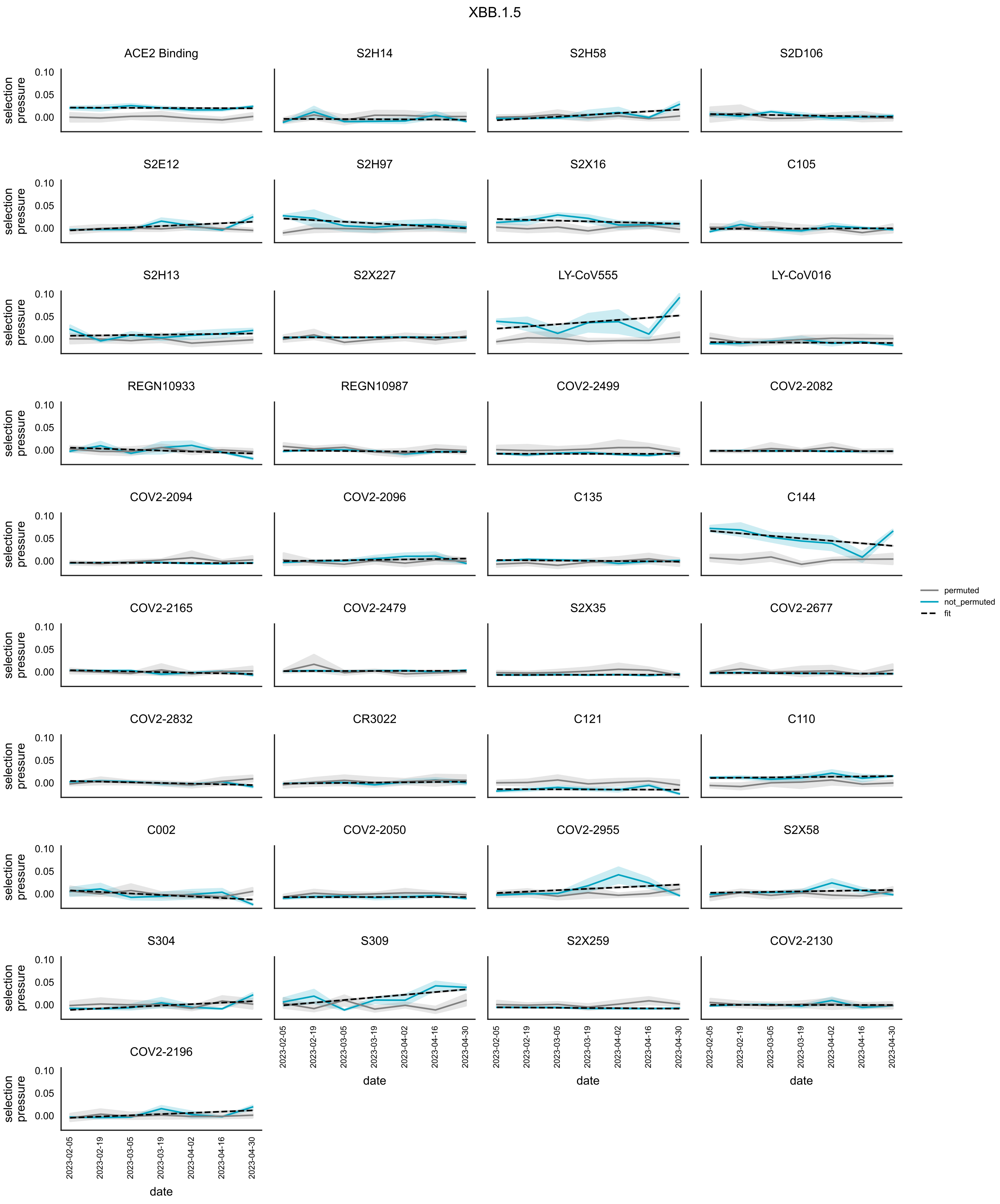


**Figure S21:** As above for the XBB.1.5 variant.

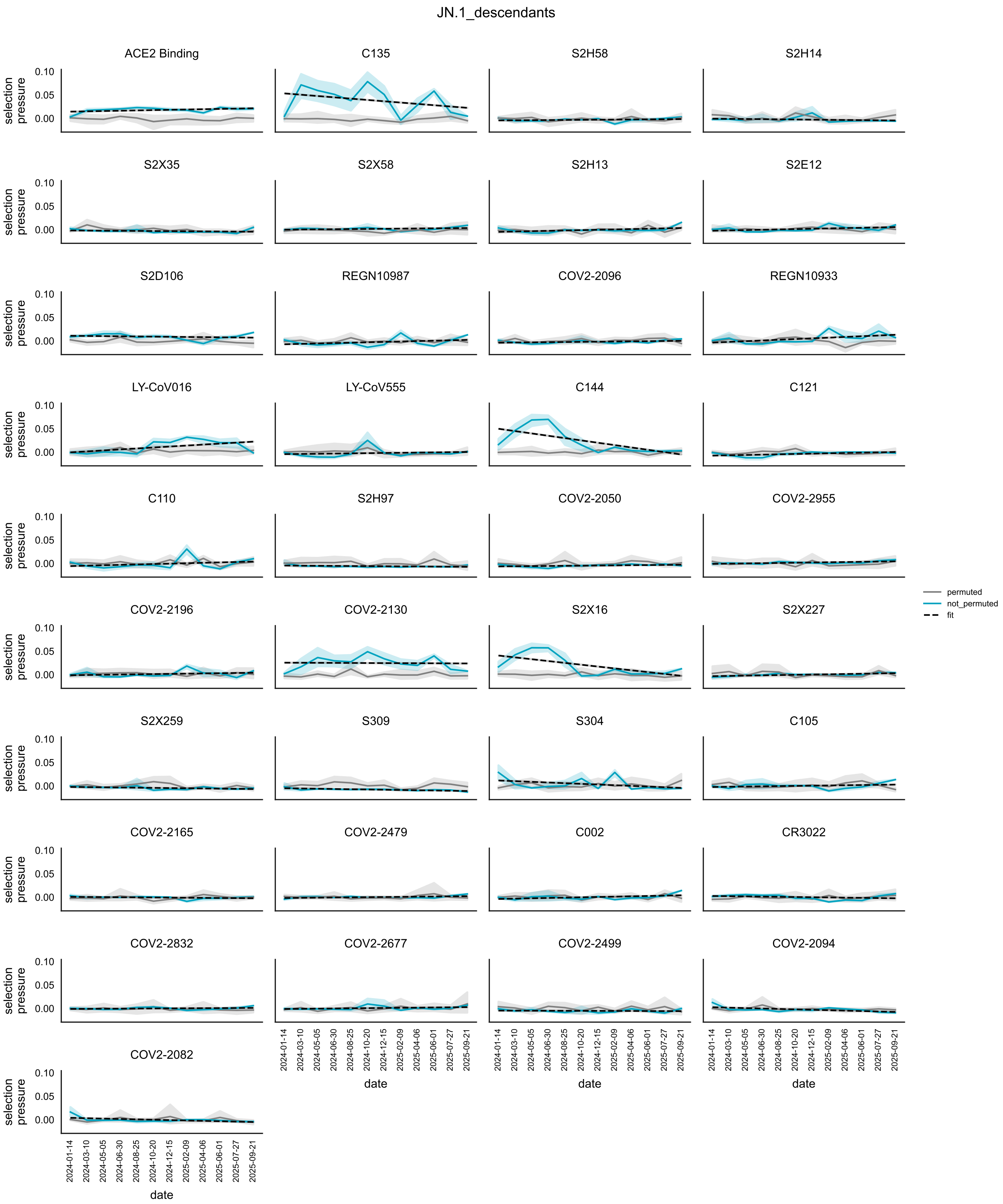


**Figure S22:** As above for the JN.1 variant and direct descendants.

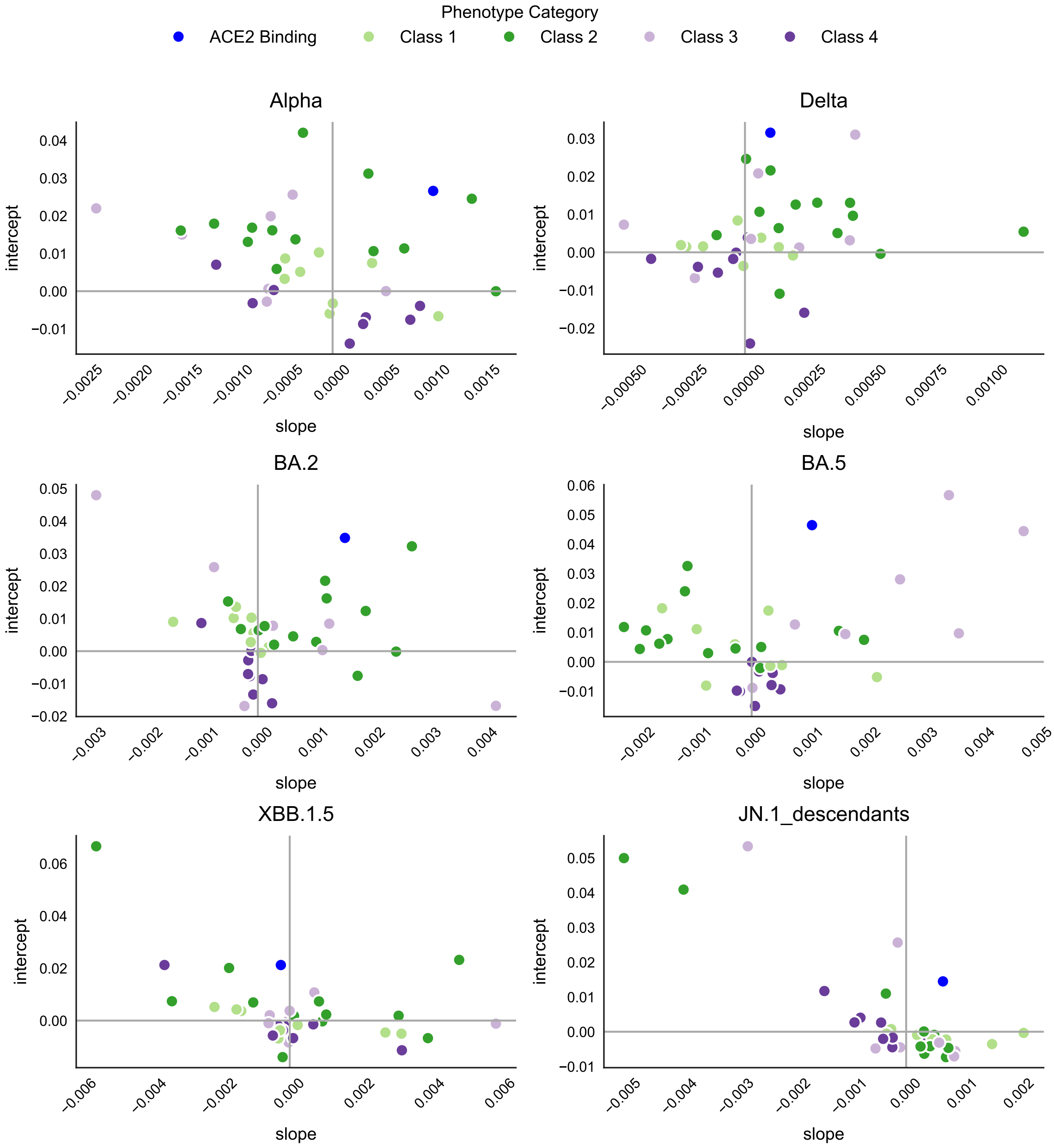


**Figure S23:** The relationship between the intercept and slope of the fitted straight line to the time-varying selection pressure trajectories for different phenotypes, within different major variants/Nextstrain clades. Datapoints are coloured according to the phenotypes category (either ACE2 binding or escape from one of the four Barnes antibody classes).

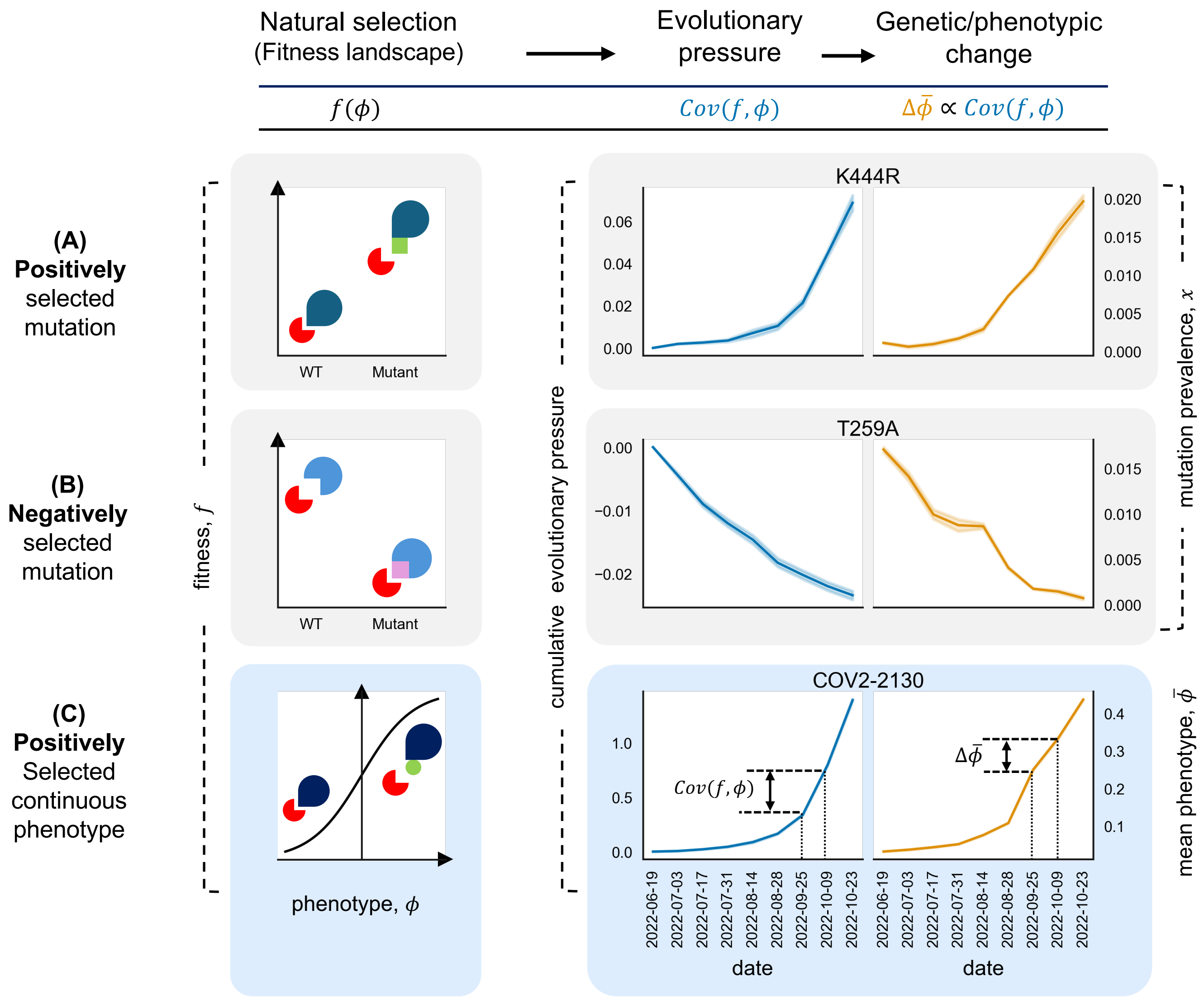


**Figure S24:** Natural selection (cause) results in genetic/phenotypic change (effect), mediated by population composition. (A, B) Left: fitness landscapes map a genotype or phenotype, , to the viral fitness f. In this example, we consider a simple binary genotype where the two genotypes (WT and mutant) differ by one mutation. The biophysical origin of viral fitness lies in the ability of a virus protein (dark blue) to escape the binding of a particular antibody (red). We show fitness landscapes for two possible mutations. One beneficial mutation (A) has a positive fitness effect $f > 0$, where the fitness effect is defined as the difference between the fitnesses of the mutant and wildtype viruses, $f = f_{mutant} - f_{WT}$, as calculated within the Omicron BA.5 variant (Nextstrain clade 22B). One deleterious mutation (B) has a negative fitness effect, $f < 0$. Middle: fitness differences within a population result in an evolutionary pressure, defined as the covariance between fitness and a phenotype, $Cov(f, \phi)$, where here the phenotype is simply the presence of the particular mutation. The evolutionary pressure depends on the fitness landscape and the genetic composition of the population. For beneficial and deleterious mutation, the evolutionary pressure is often positive and negative respectively, as in these examples. Right: the beneficial mutation (positive fitness effect, i.e. reduced binding in this example) is positively selected, leading to positive evolutionary pressure accumulating in time (blue) and driving an increase in the mutation prevalence (orange). The deleterious mutation shows the opposite trend. (C) For a continuous phenotype , viral fitness $f$ is a continuous function of the phenotype – the fitness landscape (left). This is the main object of interest in this paper. In this example, there is a positive correlation between the phenotype and the fitness. Within Omicron BA.5, the phenotype is indeed positively selected, and the estimated population-level covariance between fitness and phenotype is positive. Both the cumulative covariance (left) and the average value of the phenotype over viral sequences (right) increase proportionally with time. These dynamics approximately satisfy the Price equation: the change in the average phenotype $\Delta\bar{\phi}$ , is proportional to the evolutionary pressure, $Cov(f, \phi)$).

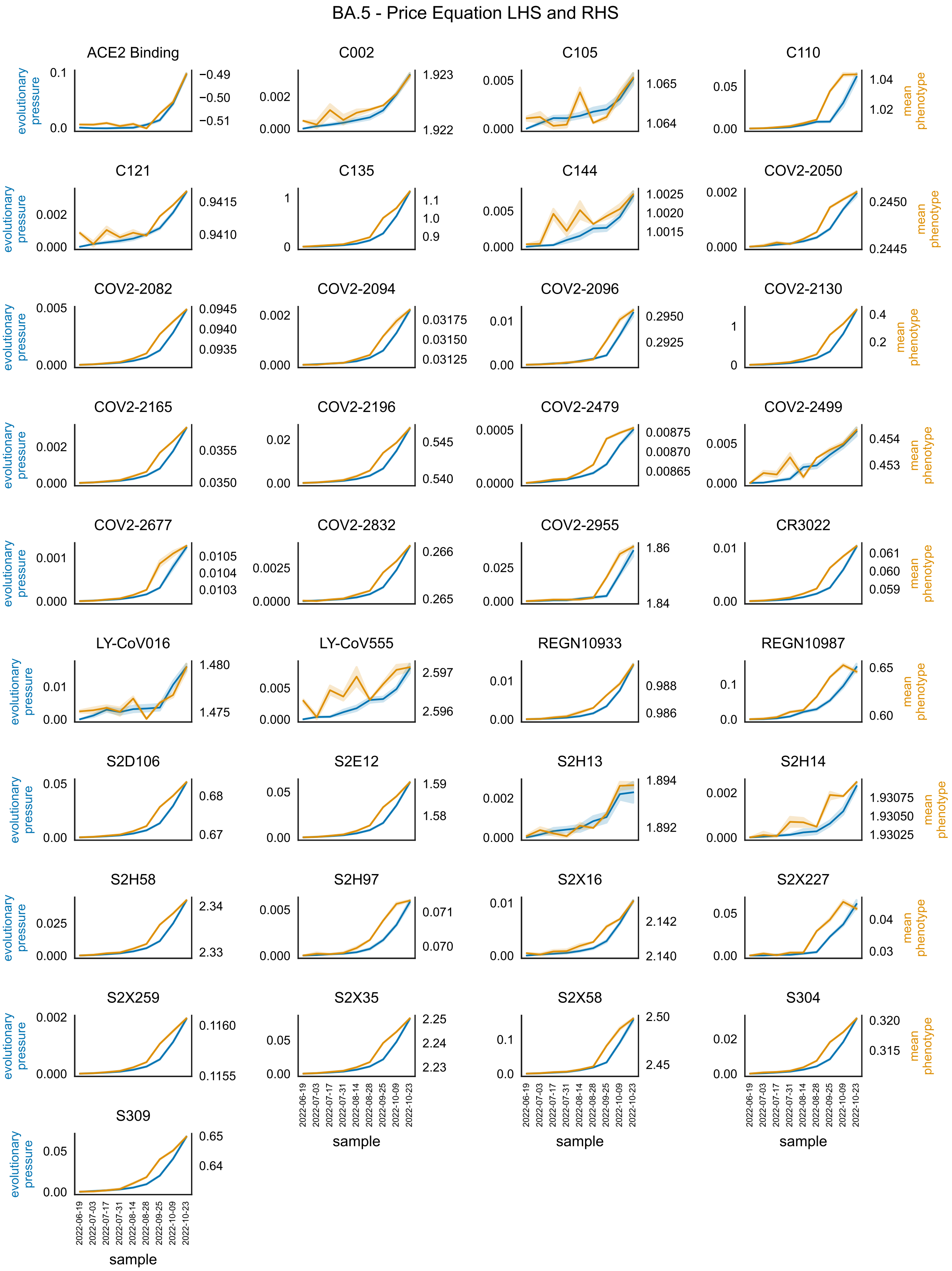


**Figure S25:** Examining the validity of the Price equation for within-clade evolution within an example Nextstrain clade, 22B (BA.5), for all phenotypes considered. On the left-hand-side (LHS, blue) is plotted at each time $t_n$, the cumulative evolutionary pressure up to that time $t_i$, for a phenotype $\phi$, i.e. $\sum_{j=1}^{i} Cov(f(t_j), \phi(t_j))$. On the RHS (orange) is plotted the mean phenotype across sequences at that time point, $\bar{\phi}(t_i)$. The two lines are approximately proportional for each phenotype, confirming the approximate validity of Price's equation: $\Delta\bar{\phi} \propto Cov(f, \phi)$.

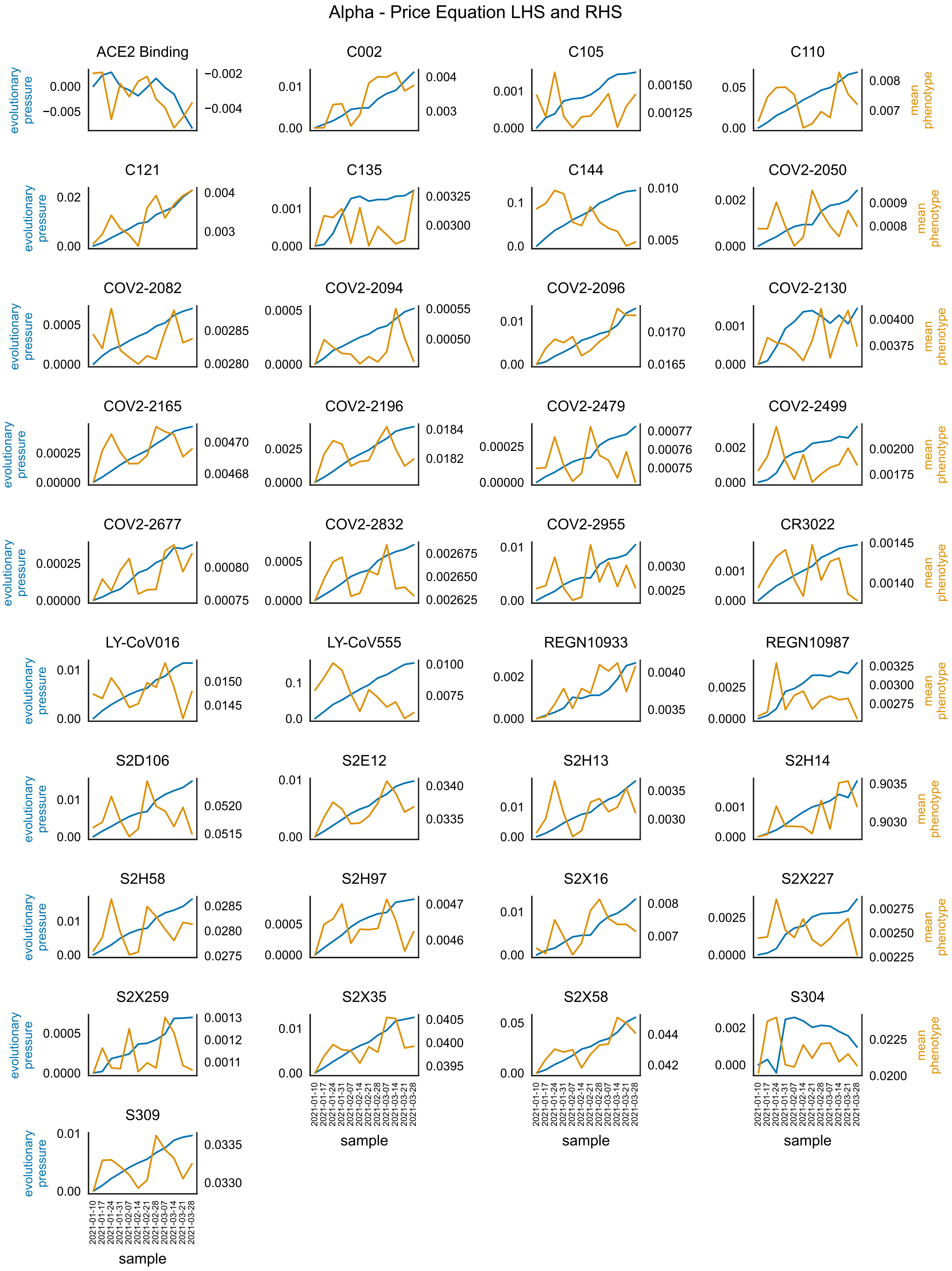


**Figure S26:** As above, for the Alpha variant. Here, however, the results for only one downsample have been plotted.

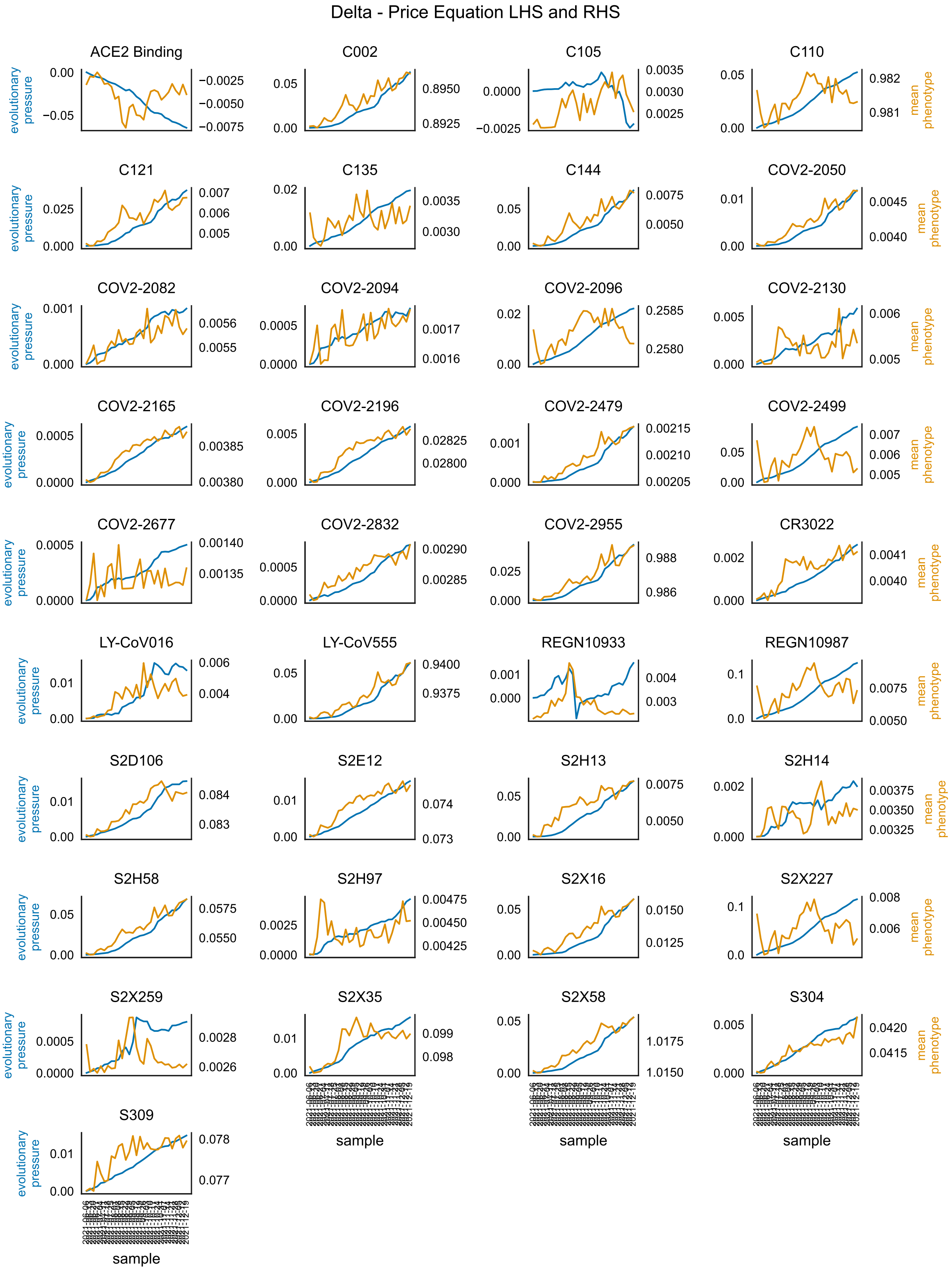


**Figure S27:** As above, for the Delta variant.

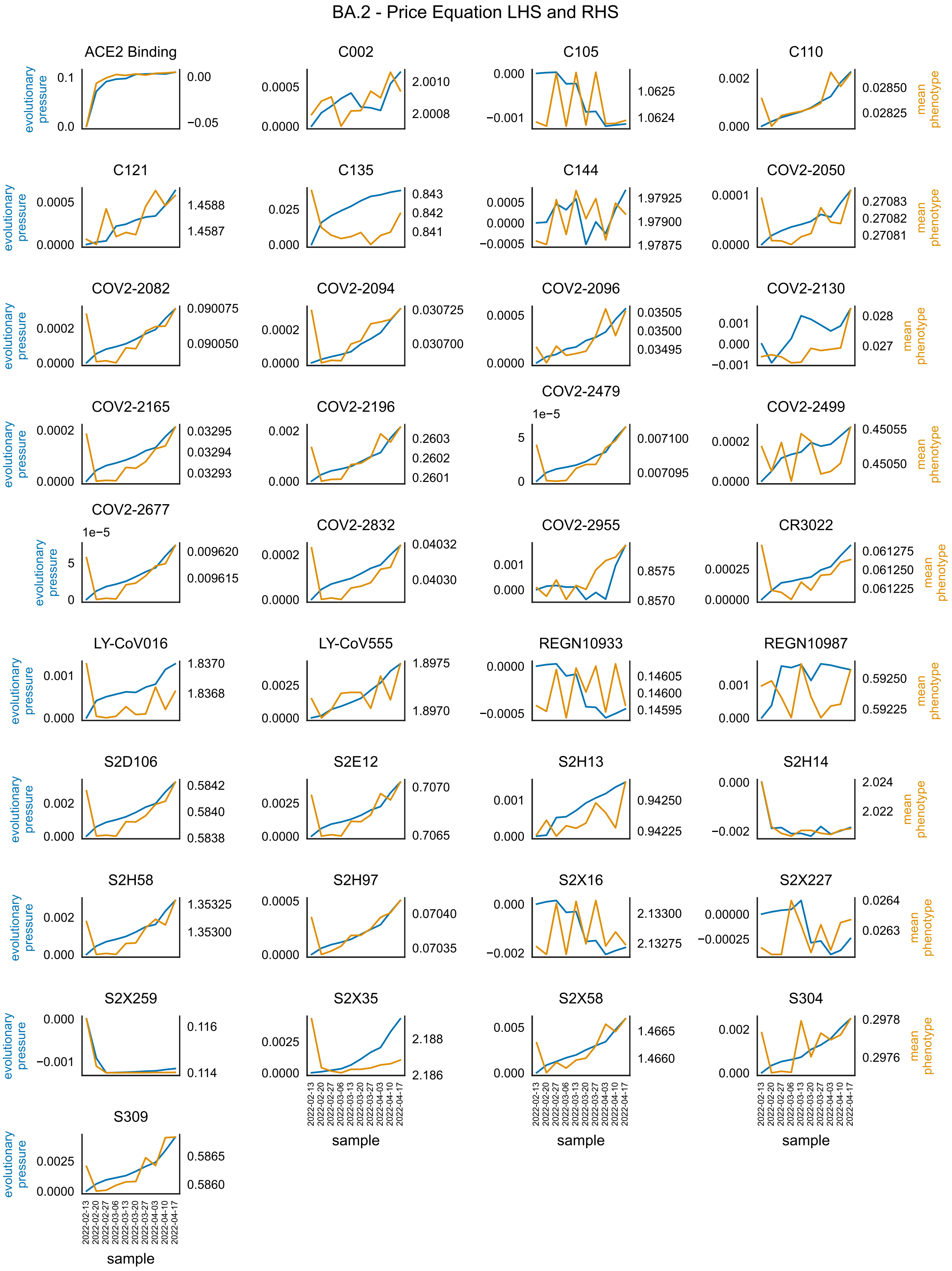


**Figure S28:** As above, for the Omicron BA.2 variant.

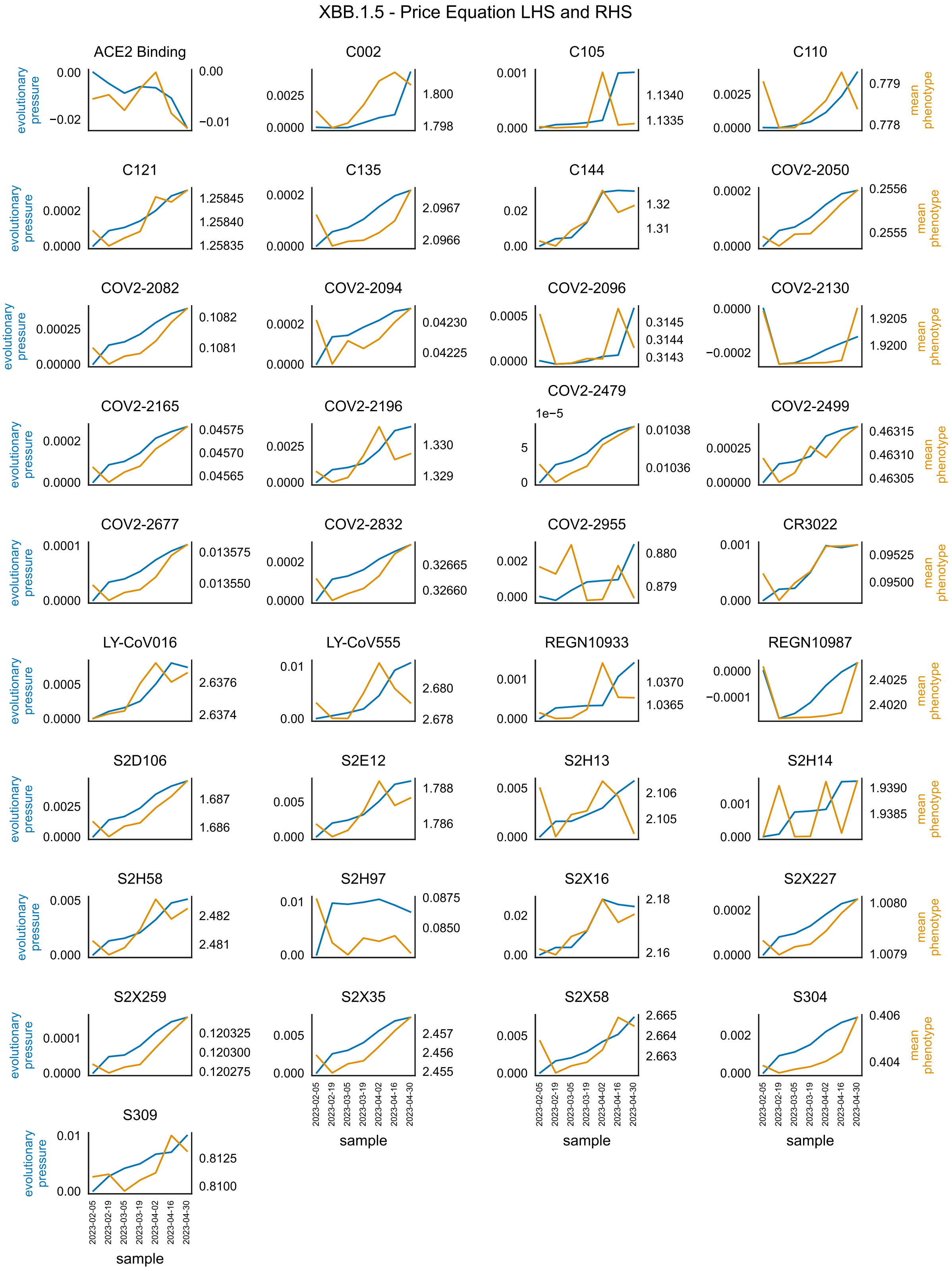


**Figure S29:** As above, for the Omicron XBB.1.5 variant.
